# Interannual temporal patterns of DeSoto Canyon macrofauna and evaluation of influence from the Deepwater Horizon

**DOI:** 10.1101/2021.04.11.438379

**Authors:** Arvind K. Shantharam, Amy R. Baco

**Affiliations:** Florida State University, Department of Earth, Ocean, and Atmospheric Sciences, 1011 Academic Way, Tallahassee, FL 32306

## Abstract

Submarine canyons are highly dynamic and productive ecosystems, but time-series studies of metazoan benthic communities in canyons are scarce. Deep-sea macrofauna from the DeSoto Canyon in the northern Gulf of Mexico were sampled annually from 2012 through 2014 from five stations within the Canyon and from two stations in 2013 and 2014 on the adjacent open slope, for analysis of interannual dynamics, temporal variability, and potential influence of the Deepwater Horizon oil spill (DwH), which occurred nearby in 2010. At a few sites, elevated abundance was observed at the start of the time-series for overall macrofauna and for deposit feeder abundance. However, diversity metrics showed no difference within stations among time points. Community and feeding guild structure varied by station, as expected, but showed no statistical difference among time points within a station. Some temporal variability was visible in temporal trajectory overlays. Cluster analyses showed grouping more by station than by time point. Metrics utilized for measuring potential oil contamination impact and overall community stress including the benthic polychaete/amphipod ratio, feeding guild abundance, macrofaunal indicators designed from the DwH, and community dispersion, generally exhibited a paucity of evidence of impact, both yearly and with site-to-site comparisons. This suggests low levels of impact in the canyon consistent with the low deposition of hydrocarbons, the timing of sampling, and quick recovery of canyon foraminifera. Taken together these results suggest relatively low levels of temporal variability within the DeSoto Canyon macrofauna with little evidence of oil influence on these sites within the studied time range.

## I. Introduction

Despite its seemingly quiescent nature, closer inspection of the deep-sea indicates natural temporal variability that can be linked to seasonal and stochastic events, e.g. surface primary productivity changes, climate, sediment slumping, and sediment resuspension from benthic storms (reviewed by Smith et al. 2009; Glover et al. 2010). Not surprisingly, natural variations in deep-sea fauna through time have also been tied to this environmental variability, with fluctuations in abundance and diversity as well as species composition (Glover et al., 2010 and references therein; Laguionie-Marchais et al., 2013; Ramalho et al., 2014; Rogers 2015). Ecological processes such as succession, disturbance, and recovery can also contribute to fluctuations in abundance and community structure in deep-sea benthic communities over time (Smith 1986; Kukert and Smith 1992; Young and Richardson 1998; Smith et al. 2002; Bernardino et al. 2010).

From a basic science perspective, since the deep sea varies through time in both environmental and faunal patterns, characterizing the range of these natural variations is critical to our understanding of deep-sea ecology. In a more applied context, quantifying the natural range of variation in deep benthic populations through time also provides a baseline for assessing natural and anthropogenic disturbances. Once the normal variation of a system has been quantified, this data can act as a baseline to assess whether any observed changes can be attributed to a disturbance event or to natural variability. However, in the most recent review of deep-sea time-series studies, Glover et al. (2010) found there were only 11 deep-sea sedimented ecosystems with multi-year time-series studies, of which only 2 were longer than 5 years. Thus, the paucity of time-series studies in the deep ocean makes it hard to determine whether changes observed after a human impact are caused by the impact, or a part of the natural variation of the system. The lack of data also leads to an inability to predict whether the effects from disturbances will be prolonged or ephemeral.

Increasing human-induced, large-scale disturbances have accentuated the imperative for more time-series studies to allow comparisons and prediction of impact outcomes (Glover and Smith 2003; Ramirez-Llodra et al. 2011; Mengerink et al. 2014). Highly dynamic, ecologically important environments such as submarine canyons in particular, need more focused studies as they are noted to be hotspots of diversity, with large spatial and biogeochemical heterogeneity. In the last decade, canyons have increasingly become targets for bottom-trawling fisheries, are exploited for oil and gas exploration, are sinks for litter and chemical pollution, and the biota inhabiting them may be sensitive to the effects of climate change over time (Fernandez-Arcaya et al. 2017; De Leo and Puig 2018).

The DeSoto Canyon is a major topographic feature of the northern Gulf of Mexico (NGOM). Environmentally, it stands as a noted sedimentary transition zone (Antoine and Bryant 1968), which can accumulate high amounts of organic matter (Morse and Beazley 2008; Silva 2017; Wei and Rowe 2019), and potentially has a strong hydrodynamic regime given its inherent topography (Clarke and Van Gorder 2016). Ecologically, it is recognized as a hotspot of meiofaunal and macrofaunal benthic abundance (Baguley et al. 2006; Wei et al. 2010, Shantharam et al in revision), benthic diversity (Wicksten and Packard, 2005), and biomass (Wei et al. 2012). To date, however, the temporal dynamics of resident macrofauna in this canyon have not been characterized, leaving a gap in baseline knowledge on how biota of this important bathymetric feature behaves ecologically through time. It has been shown that the larger NGOM macrobenthos seems subject to some interannual variability in abundance (Montagna et al. 2020) and community structure (Salcedo et al. 2016). Thus, the primary goal of this study was to test the null hypothesis that macrofauna in more localized ecosystems like the DeSoto Canyon exhibit interannual variability similar to non-canyon sites and that individual locations exhibit similar inter-year patterns.

Given the proximity of the DeSoto Canyon to the Deepwater Horizon oil spill (DwH), it is necessary to also consider whether any of the study sites were influenced by the spill. DwH was the largest oil spill in United States history and the first in the deep sea (wellhead depth 1500m). The infaunal impact in the area immediately around the DwH wellhead was extensive and impacts have been long-lasting (Montagna et al. 2013; Baguley et al. 2015; Montagna et al. 2016; Reuscher et al. 2017; Washburn et al. 2017). Further away, the picture is less clear. In the DeSoto Canyon, 40 – 185 km from the wellhead, rapidly deposited labile and non-labile hydrocarbon sedimentation was confirmed in sites throughout the canyon (Brooks et al. 2015). Impacts to sediment ecosystems included changes to the redoxcline, similar to an influx of enriched organic matter, from 2010 – 2013 (Brooks et al. 2015; Hastings et al. 2015). Oil-degrading bacteria in sediments near the deeper plumes spiked in September/October 2010 and in the summer seasons of 2012 – 2014 (Mason et al. 2014; Overholt 2018). Impacts to local benthic meiofauna were inferred from the drop in foraminiferan density and diversity, and overall sediment bioturbation (Schwing et al. 2013; Brooks et al. 2015; Schwing et al. 2015; Schwing et al. 2017).

By 2014 – 2016, the impacts to DeSoto Canyon ecosystems may have subsided. Perturbations to phytoplankton productivity at the initiation of the DwH mostly abated by 2014 and 2015 (Li et al. 2019). Signs of the mass deposition event in the canyon had ended by 2013 - 2016 (Larson et al. 2018). Bacterial communities in canyon sediment may have returned to baseline conditions by 2015 (Yang et al. 2016; Liu et al. 2017; Overholt et al. 2019) as bioturbation recommenced between 2013 – 2016 (Larson et al. 2018) and the redoxcline resumed steady-state conditions (Hastings et al. 2020). Additionally, canyon foraminiferan diversity rose and steadied by 2012 (Schwing et al. 2017; Schwing et al. 2018; Schwing and Machain-Castillo 2020). However, foraminiferans only comprise a small portion of the benthic fauna occupying this soft sediment ecosystem, leaving many facets of metazoan abundance, diversity and structure unexplored. Thus, with macrofauna sampled during the summer of 2012 – 2014 in the DeSoto Canyon, and adjacent continental slope, a further goal of this study was to test whether any observed temporal variability might be attributable to influence from the oil spill and to compare observations between communities in the canyon to those on the non-canyon slope.

## II. Materials and methods

### 2.1 Study location

The DeSoto Canyon is an S-shaped canyon located on the edge of the NW Florida outer continental shelf, cutting from the shelf to the abyssal region of the Gulf of Mexico (GOM). It is a sedimentary ecotone with siliclastic clays to the west and biogenic carbonates to the east (Gould and Stewart, 1955; Doyle and Sparks, 1980). Sediment accumulates at a rate of ~17 cm/ky (Emiliani et al., 1975) in the northwest and ~10 cm/ky in the southeast (Emiliani et al., 1975; Nürnberg et al., 2008) of the canyon. In the summer season, the head of the canyon is occupied by low salinity, biologically productive waters, driven there by cyclonic and anticyclonic eddies (Müller-Karger et al., 1991; Belabbassi et al., 2005; Walker et al., 2005; Biggs et al., 2008; Jochens and DiMarco, 2008). Strong thermohaline stratification prevents further intrusion, leaving highly oxygenated water typical of the North Atlantic Deepwater (NADW) to be the predominant water mass in the deeper (>1000 m) reaches of the canyon (Rivas et al., 2005; Morse and Beazley, 2008). Further detail on habitat characteristics are reviewed in Shantharam et al (in revision).

### 2.2 Relation to the Deepwater Horizon deposition

The DeSoto Canyon is located ~40 – 185 km east/southeast of the DwH wellhead (Fig 1) and towards the eastern edge of the total estimated 24,000 km^2^ depositional footprint of the oil spill (Chanton et al. 2014). The extent of the surface slick in relation to the canyon is depicted in Figure 1A. Plumes from the DwH spill were detected within the DeSoto Canyon at ~400, ~1000, and 1400 m depths in 2010 (Hollander et al. 2012). Within a year after the spill, rapid deposition of soluble and insoluble hydrocarbons was observed. Sediment hydrocarbon concentrations increased five times more than pre-spill levels (Romero et al. 2015; Romero et al. 2017). The distribution of radiocarbon (^14^C) of DeSoto Canyon seafloor sediments is presented in Fig 1B, more depleted values are indicative of petrocarbon from the DwH. These mostly comprised degraded, high molecular weight compounds n-alkanes (67%), low molecular weight n-alkanes (9%) and low weight PAHs (6%). This composition remained relatively unchanged for 3 years though large reductions in concentrations did occur for homohopanes (~67%) and low weight compounds (n-alkanes and PAHs, ~65% and ~66% respectively) and to a lesser degree high molecular weight n-alkanes (~43%) and PAHs (~12%) (Romero et al. 2020). Most of the deposition of this material seems to be concentrated west of the Canyon, but a few patches are observed in Fig 1B within the canyon.

**Figure 1.**
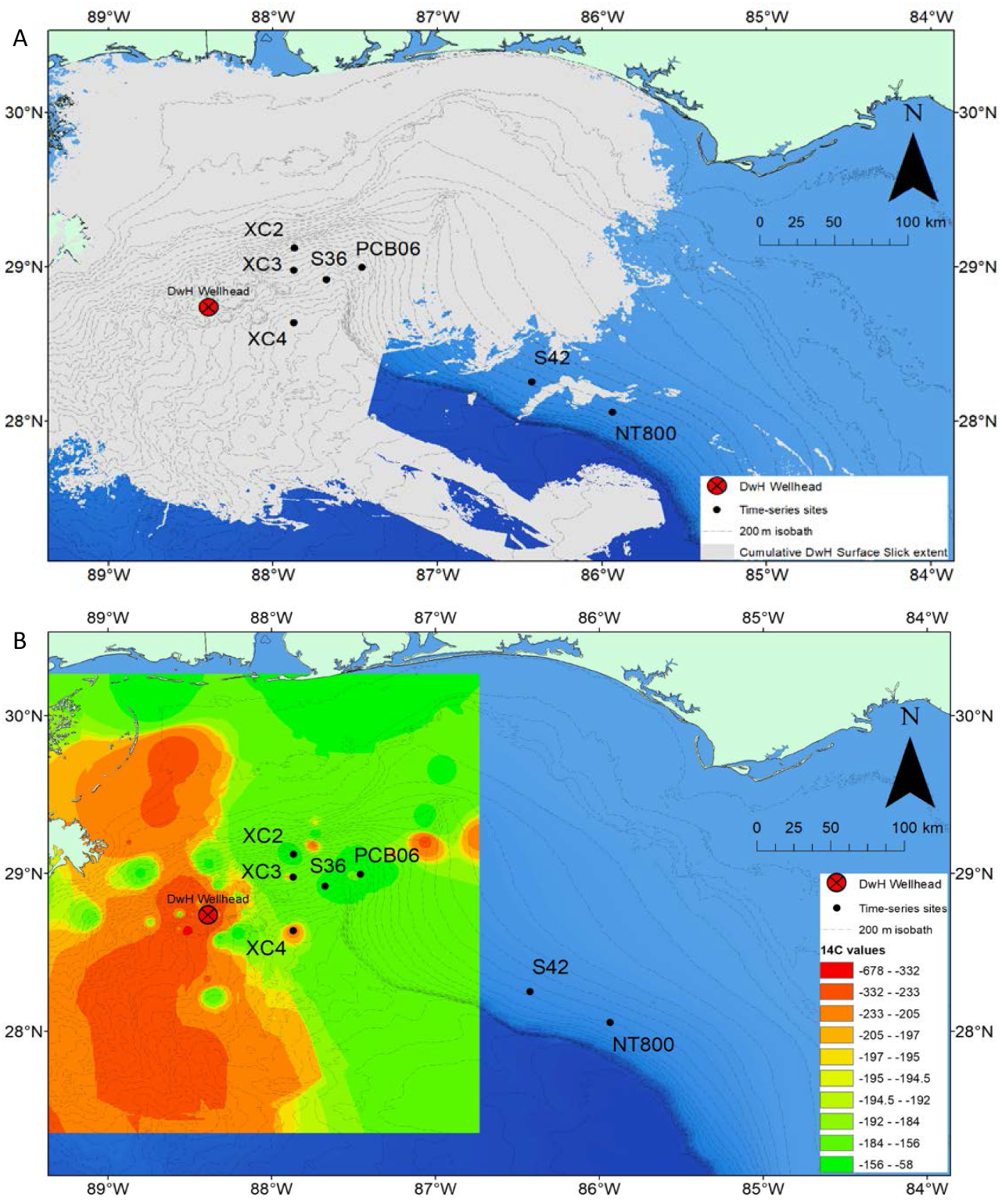
A) Map of DeSoto Canyon and open slope time-series sites. Sites were sampled from 2012 – 2014 in the DeSoto Canyon and 2013 – 2014 on the adjacent slope. The light gray overlay is the maximum extent of the surface petroleum slick (ERMA, 2019). B) Time-series sites in relation to sediment radiocarbon contoured with the Inverse Distance Weighting method with data from Chanton et al. (2014).

### 2.2 Sampling

Samples were collected during the *R/V Weatherbird II* cruise numbers WB1305, WB1306, WB1405, and WB1411. Sampling in 2012 and 2013 took place in late August to early September and sampling in 2014 from late June to early July. Five sites within the DeSoto Canyon, PCB06, XC2, XC3, S36, and XC4 were sampled (Figure 1). Three sites, PCB06, S36, and XC4, were sampled annually within the study period and XC2 and XC3 were only sampled in 2012 and 2014 (due to weather in 2013 and equipment malfunctions) (Table 1). Two sites on the adjacent eastern non-canyon slope, expected to be fully outside the oil influence, were also sampled in 2013 and 2014: S42 and NT800 (Table 1, Fig 1). Only site XC4 shows potential spill influence based on the Chanton et al. (2014) data (Fig 1B). However, all canyon sites fall under the surface footprint of the oil slick. XC2, PCB06, and XC3 of this study were also shown to have had direct deposition in 2010 (XC2 and PCB06) and 2011 (XC3) (Romero et al. 2015).

**Table 1.** Summary of time-series sampling Sites in the DeSoto Canyon and the adjacent non-canyon continental slope. An “X” indicates samples were collected in that year at that station.

| Site | Latitude | Longitude | Depth (m) | Time Points |  |  |
| --- | --- | --- | --- | --- | --- | --- |
|  |  |  |  | 2012 | 2013 | 2014 |
| S42 | 28.2528 | -86.4216 | 770 | - | X | X |
| NT800 | 28.056 | -85.9335 | 800 | - | X | X |
| PCB06 | 28.99451 | -87.4574 | 1100 | X | X | X |
| XC2 | 29.1210 | -87.8655 | 1137 | X | - | X |
| XC3 | 28.9762 | -87.8683 | 1510 | X | - | X |
| S36 | 28.91851 | -87.6722 | 1834 | X | X | X |
| XC4 | 28.6365 | -87.8685 | 2200 | X | X | X |

Sediment at each site was collected using an Ocean Instruments MC-800 multicorer capable of collecting up to eight, 10-cm diameter by 70-cm long cores per deployment. Four cores from three replicate deployments at each site were processed for resident macrofauna (sensu stricto), except for NT800 in 2013 where equipment malfunctions allowed only two successful deployments. The top 10 cm of each core was sectioned into 0 – 1, 1 – 5, and 5 – 10 cm depth fractions and fixed in buffered 10% formalin. Once returned to the laboratory, macrofauna were sieved through a 300 μm sieve and stained with Rose Bengal, then preserved in 70% ethanol. Once extracted, macrofaunal specimens were identified (usually to family), and enumerated for subsequent analyses. Family level classification is generally regarded as sufficient to discern multivariate faunal patterns in marine ecosystems (Warwick 1988, Narayanaswamy et al. 2003) and in oil impact assessments (Gomez Gesteira et al., 2003).

### 2.3 Statistical analyses

For all statistical analyses, the four cores from each deployment were combined and treated as one sample, with each deployment considered a replicate for that site in that year. This study focuses on temporal variation within sites while Shantharam et al (in revision) focuses on spatial variation among sites with a larger set of sites sampled throughout the canyon and adjacent slope in 2014. Thus all statistical analyses are designed specifically to test for temporal change within a site. Standard abundance and diversity metrics (species richness and Pielou’s evenness (J’)) were computed for each sample using the DIVERSE routine in Primer v 7.0.13 (Clarke and Gorley 2015). To test for changes in trophic structure, macrofaunal taxa were assigned to feeding guilds adapted from Demopoulos et al. (2014), which included carnivore, omnivore, deposit feeder, suspension feeder, and species harboring chemosymbiotic bacteria. Abundance values were square root transformed and Bray-Curtis similarity was calculated to measure macrofaunal community and feeding guild structure using group-average clustering. These were portrayed with non-metric multidimensional scaling (NMDS) and cluster diagrams. Samples were then summed across replicates within years and ordinated with a trajectory through time overlaid to depict temporal change in community structure.

For tests of macrofaunal abundance, diversity, and community structure, a two-way Permutational Analysis of Variance (PERMANOVA) (Anderson et al. 2008) was performed with time point as a fixed factor and station as a random factor followed by pairwise comparisons.

### 2.4 Bioindicators of oil influence

Additional oil-impact faunal metrics that might provide insights into any effects of the oil spill were incorporated into the statistical analyses. The first of these is a bioindicator known as the benthic polychaete/amphipod ratio (BPA) (Andrade and Renaud 2011), which was designed for gauging benthic impact of offshore oil and gas production in shallow water. There are no thresholds or definitive measures of impact, rather it contrasts the ratio in contaminated areas to non-contaminated areas to indicate community differences due to pollution. Higher values indicate a greater impact due to the increased presence of opportunistic polychaetes and reduction of sensitive amphipods.

The second set of indicators used were a set of designations of macrofauna taxa in terms of their tolerance to or sensitivity to oil, that were developed in the context of the DwH closer to the wellhead (Washburn et al. 2016). From a known oil spill impact zone, Washburn et al. (2016), using taxa identified to the family level, designated families as “pollution tolerant” if they were significantly higher in abundance and “pollution sensitive” if significantly lower in abundance in the oil impact zone. If taxa were close to being significantly different between impact and non-impact zones and abundances were at least 50% higher in one zone versus the other, they were deemed “possibly tolerant” or “possibly sensitive”. Taxa that did not demonstrate a significant difference from background were termed “cosmopolitan”. Collectively, these groups are hereafter termed DwH macrofaunal indicators. In the setup of the current study, these were assigned to the abundance data and the ratio of these groups to the total macrofaunal abundance of a sample were determined. The results are interpreted as a measure of impact to a site within a given year.

Finally, to get a sense of the community-level impact, two measures of community variability, the relative dispersion and the Index of Multivariate Dispersion (IMD), were computed by the MVDISP routine in Primer v 7.0.13 (Clarke and Gorley 2015). Relative dispersion was computed as the average for each station within a year. Ranging from 0 to 2, lower values generally indicate lower stress (Warwick and Clarke, 1995). IMD produces a singular value that reflects the contrast of the rank of Bray-Curtis similarities in one group against the ranked similarities of another. A value of +1 signifies all similarities among samples of a group are higher than any among the contrasting group. A value of 0 and −1 infers little to no difference between groups (Warwick and Clarke, 1995). For either dispersion metric, individual sample values cannot be computed thus statistical testing cannot be applied, and inferences are instead made by empirical contrasts.

Tests of oil-impact based on macrofauna (feeding guild abundance, BPA, and DwH macrofauna indicators) were tested in a two-way PERMANOVA with site and time as fixed factors. This was followed by pairwise comparisons to identify differences between time points and between sites within time points. Significance was determined by Monte Carlo simulation when low permutations (< 100) occurred. In order to run PERMANOVA tests of abundance, BPA, and DwH indicators, values from these metrics were root-transformed and Bray-Curtis similarities were calculated. Pairwise Euclidean similarities were calculated for untransformed species richness and Pielou’s evenness for the PERMANOVA. Choice of similarity index was made as prescribed by Anderson et al. (2008).

To lower the risk of a type I error for inter-site pairwise comparisons, a Bonferroni correction of α /k where k represents the number of comparisons was applied (see Cabin and Mitchell (2000)), thus based on a critical α of < 0.05, significance was detected if α fell below 0.005 for 2012 and 2013, and 0.0024 for 2014 for pairwise comparisons. All abundance and diversity metrics were computed and PERMANOVA tests were conducted in PRIMER v 7.0.13 (Clarke and Gorley 2015). Plots of univariate metrics were made in R 3.5.3 (R Core Team 2019).

## III. Results

### 3.1 Analysis of abundance, community and feeding guild composition and analysis of structure

A total of 2455 macrofaunal individuals across 119 families and higher taxa were identified and counted. Macrofaunal proportional abundances remained relatively stable through time for each of the sites in the canyon and on the adjacent slope (Table 2). Polychaetes dominated in each community through time, with 51 – 66% of the community composition, and the highest proportions at XC3 and XC4. Tanaids were the next most abundant, with 5.3 – 15.35% of the total abundance. Tanaids had rather high proportions for NT800 in both years, and at S36 and XC4 in 2012. Aplacophorans exhibited high abundances at XC2 in both years and bivalves exhibited high abundances in both years at XC3. In the three-year sites, amphipods and the non-tanaid crustaceans generally exhibited slight increases through time. Tanaids dropped by almost 8.5% at the deepest site of XC4 between 2012 and 2013. Bivalves, aplacophorans, and other molluscs displayed more disparate patterns with sporadic increases and decreases through time with no group showing large changes. All crustacean and molluscan groups exhibited somewhat marginal decreases in proportions in 2014 compared to 2012 in XC2. Similar trends were observed at XC3 though tanaids and bivalves increased. Polychaetes increased almost 10% in 2014 versus 2012 for XC2 but annelids otherwise did not change between years for other sites.

**Table 2.** Proportions of major macrofaunal taxa across pooled replicates of DeSoto Canyon sites (PCB06, XC2, XC3, S36, and XC4) and adjacent slope sites (S42 and NT800) by year.

|  | S42 |  | NT800 |  | PCB06 |  |  | XC2 |  | XC3 |  | S36 |  |  | XC4 |  |  |
| --- | --- | --- | --- | --- | --- | --- | --- | --- | --- | --- | --- | --- | --- | --- | --- | --- | --- |
|  | 2013 | 2014 | 2013 | 2014 | 2012 | 2013 | 2014 | 2012 | 2014 | 2012 | 2014 | 2012 | 2013 | 2014 | 2012 | 2013 | 2014 |
| Amphipoda | 4.84% | 3.42% | 3.53% | 3.47% | 1.64% | 3.14% | 3.28% | 5.82% | 3.87% | 2.16% | 0.16% | 2.24% | 2.45% | 4.88% | 0.79% | 0.49% | 1.46% |
| Tanaidacea | 8.87% | 7.12% | 12.94% | 10.92% | 8.22% | 8.45% | 7.53% | 7.15% | 5.66% | 5.30% | 8.21% | 10.37% | 8.82% | 7.20% | 15.35% | 7.80% | 6.83% |
| Other Crustacea | 5.91% | 6.55% | 6.27% | 4.22% | 5.98% | 6.29% | 4.05% | 4.83% | 2.62% | 4.52% | 2.09% | 6.91% | 8.58% | 9.51% | 7.48% | 8.78% | 11.71% |
| Aplacophora | 4.30% | 5.41% | 3.53% | 3.97% | 5.53% | 1.57% | 3.47% | 11.65% | 8.29% | 1.57% | 1.29% | 1.42% | 1.96% | 0.51% | 0.00% | 0.49% | 0.98% |
| Bivalvia | 5.65% | 5.98% | 7.45% | 4.71% | 8.37% | 7.07% | 8.88% | 9.15% | 7.60% | 11.79% | 13.53% | 5.69% | 4.66% | 6.94% | 5.51% | 7.32% | 6.34% |
| Other Mollusca | 2.96% | 0.85% | 3.92% | 1.24% | 0.30% | 1.96% | 2.12% | 0.83% | 0.41% | 0.00% | 1.93% | 0.61% | 1.96% | 2.83% | 0.79% | 2.44% | 0.49% |
| Nemertea | 4.57% | 5.70% | 4.31% | 3.47% | 2.39% | 5.30% | 4.83% | 2.50% | 3.73% | 1.77% | 4.51% | 4.27% | 2.94% | 4.11% | 3.54% | 5.37% | 4.39% |
| Polychaeta | 55.91% | 57.83% | 51.37% | 59.31% | 60.69% | 60.90% | 59.85% | 54.24% | 63.81% | 65.23% | 65.54% | 63.82% | 64.71% | 59.64% | 65.75% | 65.37% | 65.37% |
| Other Annelida | 1.34% | 1.14% | 0.78% | 1.49% | 2.24% | 1.18% | 0.77% | 0.83% | 0.83% | 3.54% | 1.29% | 2.24% | 1.23% | 1.54% | 0.39% | 1.95% | 0.49% |
| Miscellaneous | 5.65% | 5.98% | 5.88% | 7.20% | 4.63% | 4.13% | 5.21% | 3.00% | 3.18% | 4.13% | 1.45% | 2.44% | 2.70% | 2.83% | 0.39% | 0.00% | 1.95% |

PERMANOVA showed time point was not significant overall for within site abundance, species richness, or evenness (Figure 2). However, abundance was higher in 2012 for PCB06 (vs 2013, t = 3.4619, **p = 0.032**; vs 2014, t = 3.0148, **p = 0.04**) and S36 (vs 2013, t = 3.5856, **p = 0.019**; vs 2014, t = 2.9593, **p = 0.038**) (Fig 2A). Mean species richness showed no significant pairwise yearly change at any sites (Fig 2B). Average evenness for XC4 and PCB06 showed a general increase across years, while XC2 stayed about level and S42 and NT800 decreased (Fig 2C).

**Figure 2.**
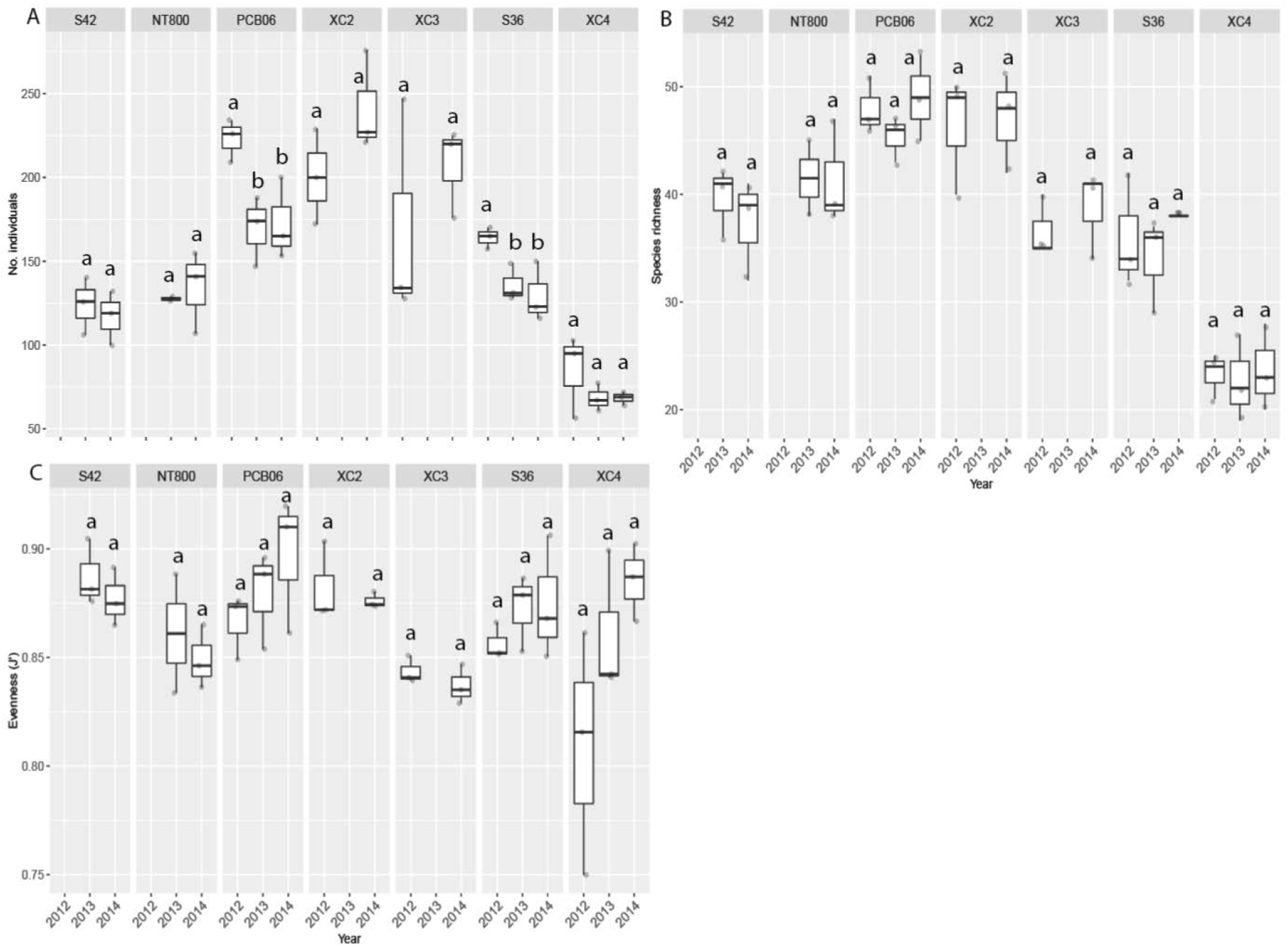
Boxplots by site and year for abundance and diversity metrics for all Sites. A) Abundance (Pseudo-F_2,49_ = 0.69131, p = 0.517). B) Species richness (Pseudo-F_2,49_ = 1.2755, p = 0.326). C) Evenness (J’) (Pseudo-F_2,49_ = 2.0279, p = 0.2). Shared letters indicate no difference.

Table 3 summarizes the feeding guild composition of the DeSoto Canyon macrofauna in 2012-2014. Deposit feeders dominated through time and across sites, representing 60% or more of all individuals, with XC4 having higher proportional abundances than any other site. Omnivores and carnivores followed in dominance and varied across stations but maintained relatively similar proportions through time. Suspension feeders and chemosymbiotic species were the least abundant, holding proportions of 9% or less across time and stations. XC3 had chemosymbiotic taxa rise almost 7% between 2012 and 2014. No consistent changes in feeding guild proportions were observed across sites except for carnivores who consistently increased through time for all canyon stations. No drastic difference in proportions were observed between slope and canyon stations.

**Table 3.** Feeding guild composition for all Sites and time points pooled across replicates.

|  | S42 |  | NT800 |  | PCB06 |  |  | XC2 |  | XC3 |  | S36 |  |  | XC4 |  |  |
| --- | --- | --- | --- | --- | --- | --- | --- | --- | --- | --- | --- | --- | --- | --- | --- | --- | --- |
|  | 2013 | 2014 | 2013 | 2014 | 2012 | 2013 | 2014 | 2012 | 2014 | 2012 | 2014 | 2012 | 2013 | 2014 | 2012 | 2013 | 2014 |
| Carnivore | 14.52% | 14.25% | 12.94% | 12.16% | 13.30% | 16.70% | 18.73% | 15.14% | 19.06% | 9.23% | 9.66% | 11.38% | 11.03% | 12.08% | 7.87% | 11.22% | 12.20% |
| Deposit | 65.05% | 65.53% | 67.06% | 66.00% | 69.36% | 64.24% | 61.58% | 68.22% | 65.88% | 65.62% | 65.70% | 63.41% | 66.91% | 62.47% | 75.59% | 70.24% | 73.17% |
| Omnivore | 14.25% | 11.40% | 11.37% | 12.90% | 7.92% | 12.38% | 9.85% | 9.65% | 8.43% | 16.50% | 11.59% | 17.48% | 14.71% | 16.97% | 7.87% | 10.73% | 7.32% |
| Suspension feeder | 3.23% | 5.13% | 5.10% | 7.69% | 6.88% | 5.11% | 6.76% | 5.32% | 5.80% | 6.29% | 4.03% | 7.32% | 6.62% | 7.46% | 7.87% | 7.32% | 5.37% |
| Chemo-symbiotic | 2.96% | 3.70% | 3.53% | 1.24% | 2.54% | 1.57% | 3.09% | 1.66% | 0.83% | 2.36% | 9.02% | 0.41% | 0.74% | 1.03% | 0.79% | 0.49% | 1.95% |

Cluster analysis of macrofaunal community structure showed samples clustered mostly by depth and by station and somewhat by year within station (Fig 3A). Samples from XC4, the deepest station at 2200 m, branched off as the most disparate cluster. Open slope stations S42 and NT800 grouped together in another single, separate cluster with a high degree of overlap in structure between the sites, regardless of time point. Among the remaining samples, two of the XC3 samples from 2012 cluster outside any other group. All time points for S36 form a cluster, and all time points from PCB06 and XC2 cluster together with the remaining XC3 samples. A NMDS of community structure (Fig 3B) largely reflected groups delineated by the cluster analysis. Tests of community structure mainly indicated a significant difference by station (**p < 0.001**, Table 4) rather than by time point (p = 0.17, Table 4). The interaction of time point and station also produced a low p-value (**p = 0.004**, Table 4). Despite these results, the trajectory plots of community structure for each station suggests some interannual variability in community structure at each site (Fig 3C). Two-year time-series sites, XC2 and XC3, showed relatively little change between 2012 and 2014. All three-year sites, PCB06, S36, and XC4, seemed to experience a larger change in structure going from 2012 to 2013, compared to 2013 to 2014.

**Figure 3.**
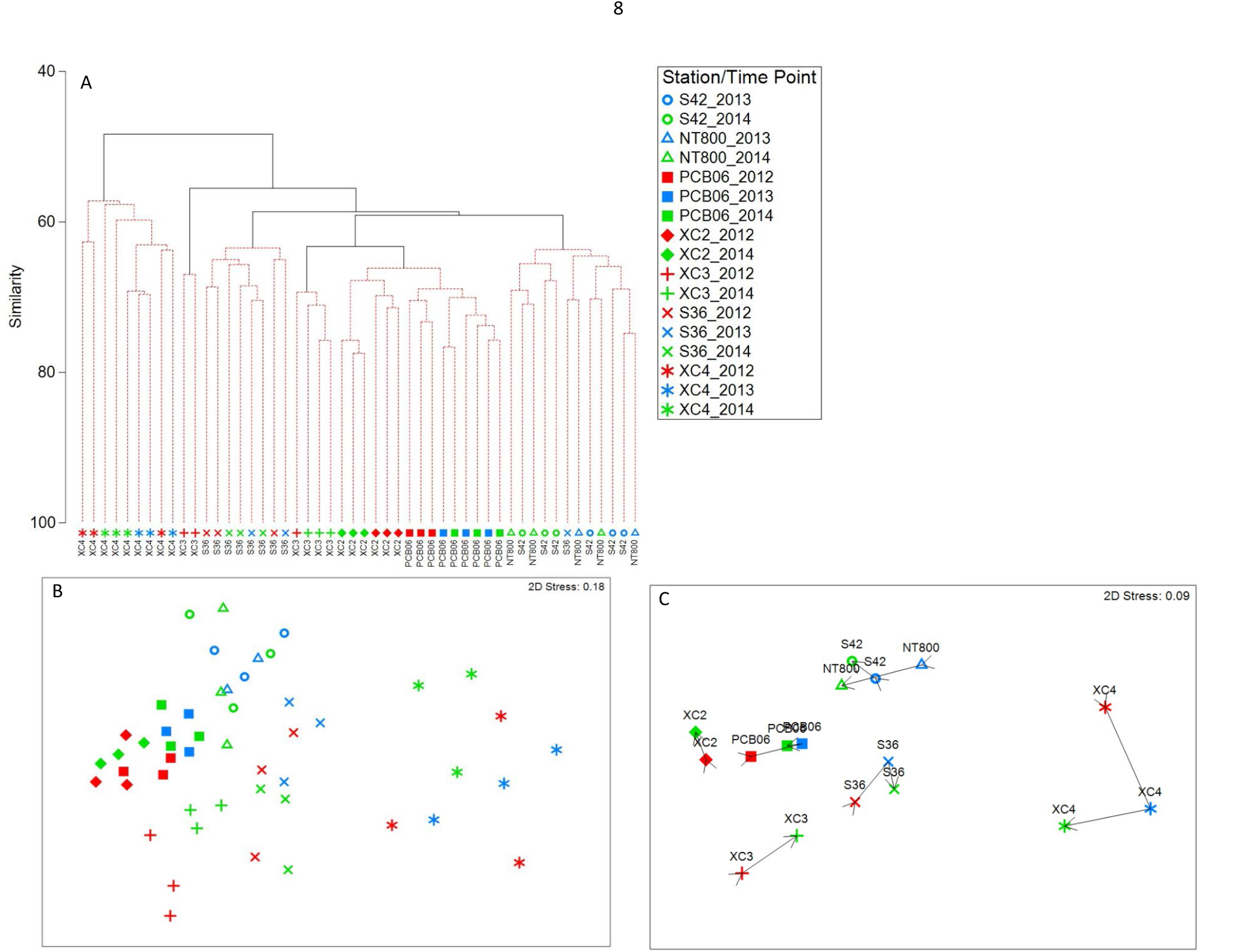
Cluster analysis based on Bray-Curtis similarities of macrofauna abundance of DeSoto Canyon time-series sites and the open slope sites from 2012 – 2014. B) Non-metric multidimensional scaling of macrofauna community structure. C) Pooled samples for each time point at each Site with trajectory overlays from 2012-2014.

**Table 4.** PERMANOVA results for the test of macrofauna community structure among and between time points from 2012 – 2014 across all sites. Bolded values indicate significant differences.

|  |  | Community Structure |  |  |  | Feeding guild Structure |  |  |  |
| --- | --- | --- | --- | --- | --- | --- | --- | --- | --- |
| Source | df | SS | Pseudo-F | P(perm) | perms | SS | Pseudo-F | P(perm) | perms |
| Time point | 2 | 1990.2 | 1.343 | 0.17 | 998 | 79.122 | 0.60557 | 0.729 | 999 |
| Site | 6 | 19489 | 5.9327 | **0.001** | 997 | 4697.8 | 15.077 | **0.001** | 999 |
| Time point x Site** | 8 | 5929.9 | 1.3539 | 0.004 | 999 | 522.77 | 1.2583 | 0.208 | 998 |
| Res | 33 | 18067 |  |  |  | 1713.7 |  |  |  |
| Total | 49 | 46737 |  |  |  | 7330 |  |  |  |

Feeding guild structure cluster analysis showed a higher percent similarity among samples but did not group as distinctly by station as the data based on family identification, or by time point, but some depth-based clustering was observed (Fig 4A). XC4 again formed its own cluster. All other stations formed a second, larger cluster that split into 2 groups, one containing 2012 and 2014 samples from PCB06, XC2 and three of the XC3 samples, the other with the remaining stations and time points. Feeding guild structure depicted by NMDS, affirmed the cluster analysis (Fig 4B). Community structure did not show a strong grouping by time point (p = 0.729, Table 4) nor by the interaction (p = 0.208, Table 4). Trajectories of structure across time illustrate stronger changes from the beginning of time-series (compared to the same graph based on families Fig 3B) for all stations except S42 (Fig 4C) that suggest modest interannual variability. Three-year time-series sites in the canyon suggest less differentiation between 2013 and 2014 (which are also the only years S42 was sampled) than compared to the differentiation of either year to the 2012 time point.

**Figure 4.**
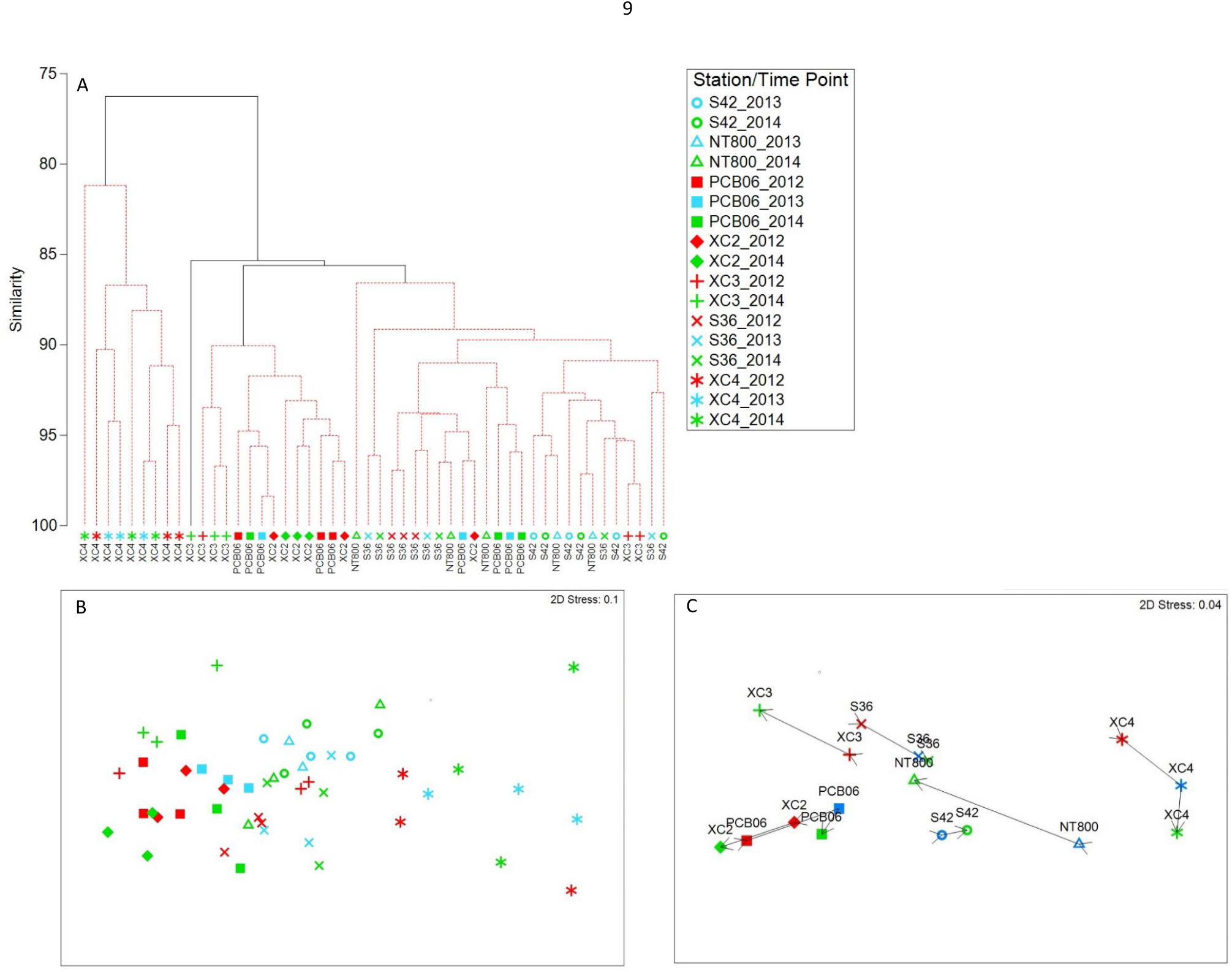
A) Cluster analysis based on Bray Curtis similarities of macrofauna feeding guilds for DeSoto Canyon timeseries sites and open slope sites from 2012 – 2014. B) Non-metric multidimensional scaling of feeding guild structure. C) Pooled samples for each time point at each Site with trajectory overlays from 2012-2014.

### 3.2 Tests of Deepwater Horizon indicators

The various indicators employed to gauge the impact of the Deepwater Horizon indicated minimal differences through time. The BPA index exhibited a high variance in 2012 across almost all sites sampled in that year (Fig 5). S36 show a gradual decline from 2012 - 2014 while XC2, NT800 and S42 seemed to show a slight increase. Due to the absence of amphipods in two of the three replicates for XC3 in 2014, the BPA could not be computed thus the whole station was excluded from analysis. Though the BPA could not be computed for XC3 in 2014, polychaete abundance for that year is plotted to contrast the BPA computed in 2012 and shows the high number of polychaetes in that year. Through time, average BPA ratios for open slope stations in 2013 (10.46 – 15.16) were generally lower than canyon stations in the same year (19.88 – 37.83) while 2014 open slope (15.00 – 15.89) and canyon (12.33 – 17.48) values were more equivalent except for XC3 where only polychaetes were recorded at 122 individuals and for XC4 with a mean BPA of 26.06. Time point was not significant for the comparison of BPA but site (**p < 0.001**, Table 5) and the interaction of site and time point were (**p = 0.021**, Table 5). The only pairwise comparisons where effects were observed were in 2013 when average BPA for S36 (20.6) (**p = 0.016**, Table 5) and XC4 (37.8) (**p = 0.013**, Table 5) were greater than S42 (10.16).

**Figure 5.**
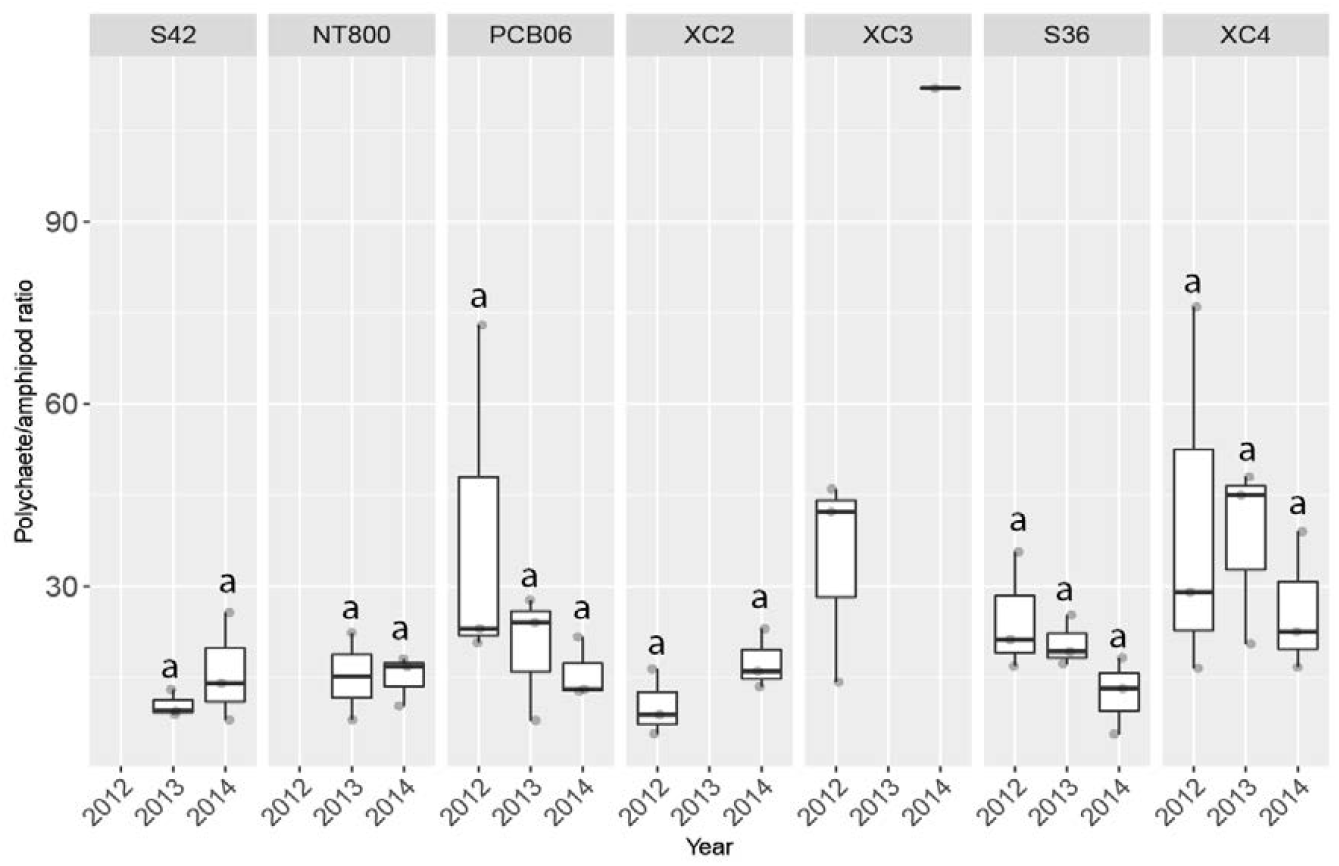
Benthic polychaete/amphipod ratio (BPA) across time for all sites (Pseudo-F_2,49_ = 1.0892, p = 0.381). No amphipods occurred in two replicates for XC3 in 2014 and therefore it was excluded from analysis but the polychaete abundance for 2014 is plotted for comparison. Shared letters indicate no difference.

**Table 5.** Summary of PERMANOVA results for the benthic polychaete/amphipod ratio. Sample size n = 3 for all groups in pairwise comparisons except for NT800 in 2012 where n = 2.

|  | df | SS | Pseudo-F | p-value |  |  |
| --- | --- | --- | --- | --- | --- | --- |
| Time Point | 2 | 986.01 | 1.0892 | 0.381 |  |  |
| Station | 6 | 8018.4 | 2.9524 | **0.001** |  |  |
| Time Point x Station | 8 | 5924.6 | 1.6361 | **0.021** |  |  |
| Residual | 33 | 14937 |  |  |  |  |
| Total | 49 | 30137 |  |  |  |  |
| Site pairwise comparisons |  |  |  |  |  |  |
|  |  | 2012 |  | 2013 |  | 2014 |
| Groups | t | P(MC) | t | P(MC) | t | P(MC) |
| S42, NT800 | - | - | 0.65053 | 0.569 | 0.16321 | 0.914 |
| S42, XC2 | - | - | - | - | 0.5 | 0.661 |
| S42, XC3 | - | - | - | - | 1.511 | 0.155 |
| NT800, XC2 | - | - | - | - | 0.65157 | 0.568 |
| NT800, XC3 | - | - | - | - | 1.5468 | 0.144 |
| XC2, XC3 | 2.2834 | 0.075 | - | - | 1.532 | 0.153 |
| PCB06, S36 | 0.70725 | 0.523 | 0.42152 | 0.702 | 0.81779 | 0.448 |
| PCB06, XC4 | 5.017E-2 | 0.993 | 1.4059 | 0.226 | 1.5176 | 0.192 |
| PCB06, XC2 | 2.2968 | 0.062 | - | - | 0.45772 | 0.662 |
| PCB06, XC3 | 0.11061 | 0.97 | - | - | 1.5417 | 0.131 |
| PCB06, S42 | - | - | 1.2834 | 0.272 | 0.24232 | 0.869 |
| PCB06, NT800 | - | - | 0.40902 | 0.709 | 0.18936 | 0.872 |
| S36, XC4 | 0.70043 | 0.511 | 1.8634 | 0.125 | 1.7718 | 0.141 |
| S36, XC2 | 2.3285 | 0.086 | - | - | 1.107 | 0.353 |
| S36, XC3 | 0.61287 | 0.56 | - | - | 1.5353 | 0.137 |
| S36, S42 | - | - | 4.0848 | **0.016** | 1.2571 | 0.677 |
| S36, NT800 | - | - | 1.0185 | 0.37 | 1.6602 | 0.533 |
| XC4, XC2 | 2.176 | 0.087 | - | - | 1.2071 | 0.303 |
| XC4, XC3 | 0.14419 | 0.929 | - | - | 1.4755 | 0.172 |
| XC4, S42 | - | - | 3.9943 | **0.013** | 1.2571 | 0.299 |
| XC4, NT800 | - | - | 1.7854 | 0.159 | 1.6602 | 0.162 |

Among feeding guild abundances, two-way PERMANOVA did not indicate differences for time points but did among stations for most feeding guilds (Pseudo-F_6,49_ = 8.4258 – 25.322, p < 0.05, Table A1) except for suspension feeders (Pseudo-F_2,49_ = 1.4517, p = 0.15) (Table A1). The interaction term was only significant for chemosymbiotic species (Pseudo-F_8,49_ = 2.3414, **p = 0.03**). A couple of pairwise site comparisons for chemosymbiotic feeders resulted in negative test statistics (Table A1), an artifact of PERMANOVA whose meaning remains elusive (Anderson et al., 2008), and thus were not interpreted. Time point pairwise tests only resulted in differences for deposit feeders at only PCB06, with larger abundances in 2012 (vs 2013, t = 2.7542, **p = 0.039**; vs 2014, t = 3.4156, **p = 0.022**) (Fig 6B) and chemosymbiotic taxa showed a significantly higher abundance at XC3 in 2014 versus 2012 (t = 2.7919, **p = 0.014**) (Fig 6E). XC2 had higher proportional abundances of carnivores in 2014, but the difference was not significant (t = 2.5478, p = 0.074).

**Figure 6.**
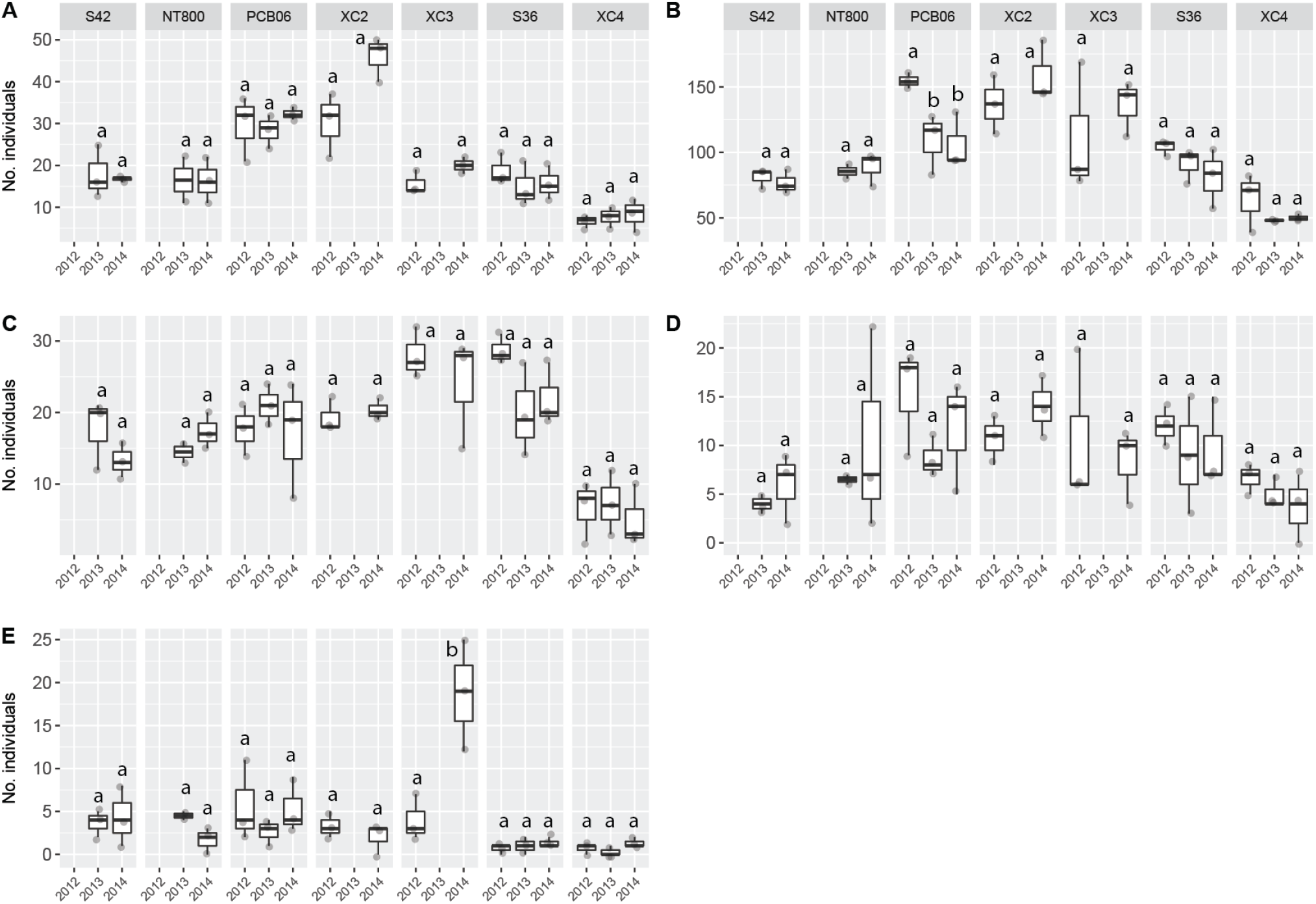
Boxplots of feeding guild abundances for all Sites by year. A) Carnivore (Pseudo-F = 0.89508, p = 0.415). B) Deposit feeders (Pseudo-F = 1.4538, p = 0.244). C) Omnivores (Pseudo-F = 0.64449, p = 0.576). D) Suspension feeders (Pseudo-F = 1.1643, p = 0.347). E) Chemosymbiotic feeders (Pseudo-F = 0.70251, p = 0.537). Shared letters indicate no differences.

Site-to-site carnivores in 2012, XC4 (mean 6.7 ind) were significantly less abundant than PCB06 (mean 29.7 ind, **p = 0.004**), XC2 (30.3 mean ind, **p = 0.001**) and S36 (mean 18.7 ind, **p = 0.004**). No differences were observed in 2013 but by 2014, the average number of carnivores at S42 (16.7 ind) was lower than PCB06 (32.3 ind, **p = 0.001**) and XC2 (46 ind, **p = 0.001**), while XC2 was greater than XC3 (20 ind, **p = 0.002**). Deposit feeders were more abundant at PCB06 than S36 (average 154.7 ind against 104 ind, **p = 0.003**) in 2012. In 2013 at S42 (80.7 mean ind) and NT800 (85.5 mean ind) (**p = 0.002**) deposit feeders were greater than XC4 (48 mean ind, **p = 0.001**). XC4 was less abundant in deposit feeders than S36 (76.7 mean ind, **p = 0.003**) in 2013. Non-canyon stations in 2013, S42 (80.7 ind) and NT800 (85.5) also showed higher abundance than XC4 (48 ind) (**p = 0.001** and **0.002** respectively). Average abundance for deposit feeders in 2014 for S42 (76.7 ind) was less than XC2 (159 ind) (**p = 0.002**). No inter-site comparisons were significant for omnivores and suspension feeders. Chemosymbiotic feeders were only significant in 2014 where XC3 (mean 18.7 ind) was higher than S36 (mean 1.3 ind) and XC4 (mean 1.3 ind) (**p=0.002** respectively). Test results of site pairwise comparisons are summarized in Table A2 of the supplementary materials.

DwH macrofaunal indicator species proportions occupied a large range of values across time points. Cosmopolitan taxa (Fig 7A) generally held high proportions, averaging about 18.5 – 35% across all sites. Tolerant taxa (Fig 7B) exhibited average proportions of 10 – 16% for non-canyon stations and 16 – 40% for sites within the canyon. Possibly tolerant taxa (Fig 7C) held low proportions, 0 – 5.4%, across all sites. Possibly sensitive proportions averaged 4.4 to 18.7% (Fig 9D) and sensitive groups averaged 17.6 – 47.5% in proportion (Fig 7E). Two-way PERMANOVA indicated a significant difference among sites (Pseudo-F_8,49_ = 5.1431 – 23.471, **p < 0.05**, Table A2), but not among time points (Table A2). The interaction term was only significant for sensitive taxa (Pseudo-F_8,49_ = 2.2515, **p = 0.027**, Table A2). Pairwise tests, however, yielded differences where global tests did not find them. XC2 and XC3 demonstrated some of the sharpest differences between years. Possibly tolerant groups were observed to be more abundant in 2014 compared to 2012 at XC2 (t = 3.5601, **p = 0.024**) and fewer in XC3 (t = 3.0391, **p = 0.045**) for the same time period. Further, S36 exhibited greater possibly sensitive group proportions in 2012 compared to 2013 (t = 3.4796, **p = 0.032**) but not versus 2014.

**Figure 7.**
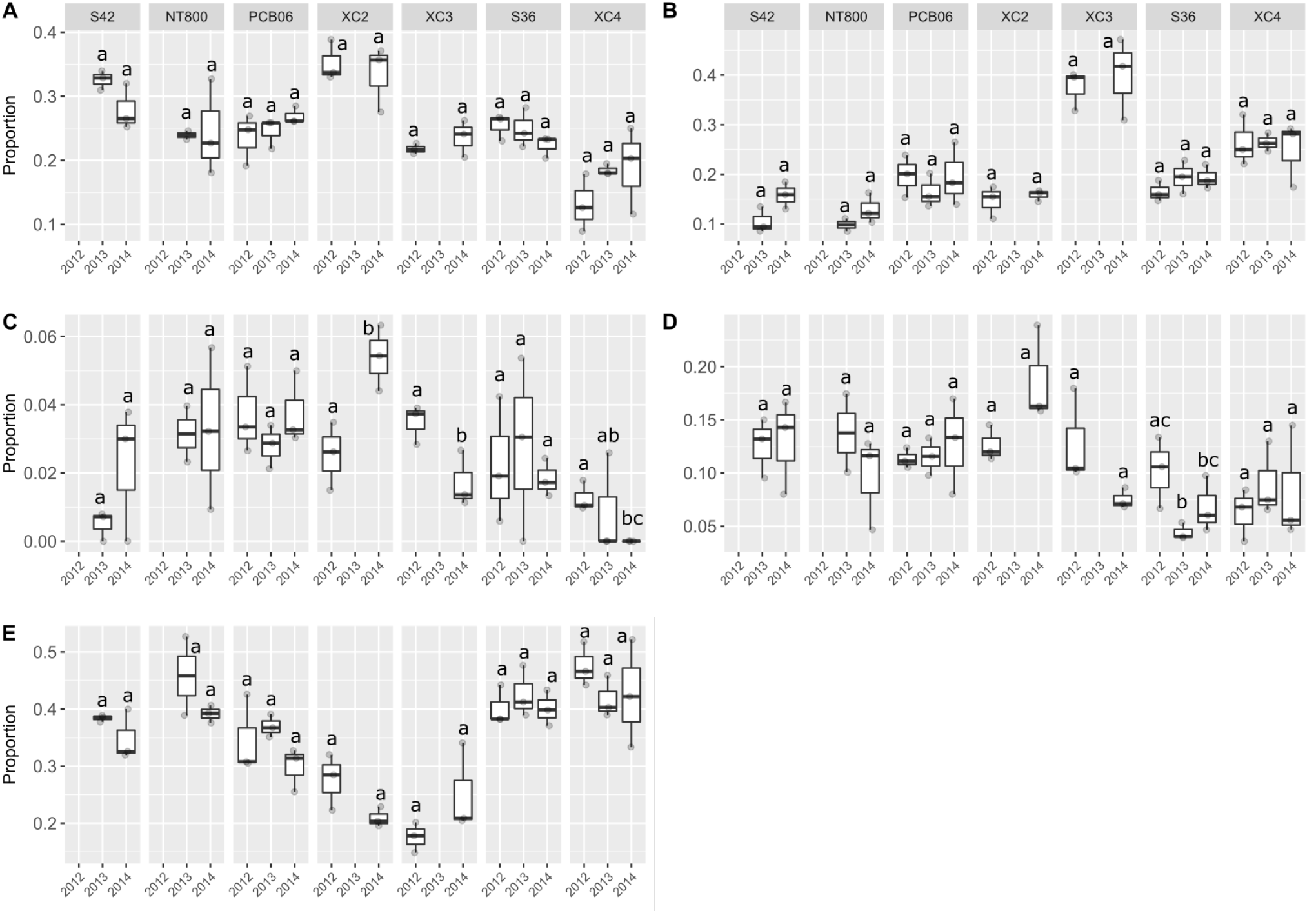
Boxplots of DwH indicator taxa community proportions following the categories of Washburn et al. (2016). A) Cosmopolitan (Pseudo-F = 0.91342, p = 0.424). B) Tolerant (Pseudo-F = 2.1075, p = 0.135). C) Possibly tolerant (Pseudo-F = 0.21704, p = 0.786). D) Possibly sensitive (Pseudo-F = 0.34496, p = 0.72). E) Sensitive (Pseudo-F = 1.5825, p = 0.217). Shared letters indicate no difference.

Site-by-site test results are summarized in Table A2. By site, cosmopolitan taxa constituted larger proportions for XC2 (mean 35.2%) than XC3 (mean 21.8%) **(p = 0.002)** in 2012. In 2013, cosmopolitan taxa proportions were higher in S42 (32.6%) than NT800 (24.0%) **(p = 0.004)**. S42 was also higher than XC4 (average 18.5%) **(p = 0.001)**. No differences were observed by 2014. Tolerant taxa proportions were only higher for XC3 (mean 37.5%) compared to S36 (24.9%) **(p = 0.001)** in 2012. In 2013 average tolerant taxa proportions at XC4 (26.4%) were lower than non-canyon stations S42 (0.5%) **(p = 0.004)** and NT800 (9.8%) **(p = 0.003)**, and by 2014 no differences were observed. Average proportions of possibly tolerant taxa were only different in 2012 where XC3 (3.5%) was greater than XC4 (1.3%) **(p = 0.003)**, and 2014 where XC2 (5.4%) was greater than XC4 (0.0%) **(p = 0.001)**. No pairwise differences were observed between sites for possibly sensitive taxa. Sensitive taxa community proportions in 2012 were higher at S36 (40.2%) and XC4 (47.5%) **(p = 0.001)** compared to XC3 (17.6%) **(p = 0.001)**. No differences were observed in 2013, but by 2014 proportions of sensitive taxa were higher at NT800 (average 39.2%) **(p = 0.001)** and S36 (40.1%) **(p = 0.002)** than XC2 (21.0%).

Community dispersion ranged from 0.12 – 1.693 among stations and years (Table 6). Non-canyon slope sites had dispersion values ranging from 0.87-1.24. PCB06 and XC2 had some of the lowest values of any station, ranging from 0.12 – 0.733, indicating generally low community stress. XC3 exhibited a decrease with a dispersion value of 1.28 in 2012 and 0.413 in 2014. S36 had relatively high values (1.24 – 1.52), but XC4 exhibited the highest dispersion, ranging from 1.693 to 1.8.

**Table 6.** Relative multivariate dispersion of macrofaunal communities at each site by year.

| Site | 2012 | 2013 | 2014 |
| --- | --- | --- | --- |
| S42 | - | 0.867 | 1.053 |
| NT800 | - | 1.24 | 1.2 |
| PCB06 | 0.533 | 0.373 | 0.733 |
| S36 | 1.173 | 1.52 | 1.24 |
| XC2 | 0.627 | - | 0.12 |
| XC3 | 1.28 | - | 0.413 |
| XC4 | 1.8 | 1.293 | 1.693 |
| Pairwise comparisons | 2012 IMD | 2013 IMD | 2014 IMD |
| S42, NT800 | - | -1 | -0.111 |
| S42, XC2 | - | - | 1 |
| S42, XC3 | - | - | 0.778 |
| NT800, XC2 | - | - | 1 |
| NT800, XC3 | - | - | 0.556 |
| XC2, XC3 | -1 |  | -0.556 |
| PCB06, S36 | -0.778 | -1 | -0.778 |
| PCB06, XC4 | -1 | -1 | -1 |
| PCB06, XC2 | -0.556 | - | 1 |
| PCB06, XC3 | -1 | - | 0.556 |
| PCB06, S42 | - | -0.778 | -0.556 |
| PCB06, NT800 | - | -1 | -0.556 |
| S36, XC4 | -1 | 0.111 | -0.778 |
| S36, XC2 | 0.778 | - | 1 |
| S36, XC3 | -0.111 | - | 0.778 |
| S36, S42 | - | 1 | 0.333 |
| S36, NT800 | - | 0.333 | -0.111 |
| XC4, XC2 | 1 | - | 1 |
| XC4, XC3 | 0.778 | - | 1 |
| XC4, S42 | - | 0.556 | 1 |
| XC4, NT800 | - | 0.333 | 0.333 |

Pairwise IMD for the 2012 inter-site comparisons generally indicated that dispersion was only different for S36 compared to XC2 and XC3 vs XC4 (Table 6). In 2013, S36 and XC4 showed some slight differentiation in dispersion and stronger differentiation compared to non-canyon stations S42 and NT800 and to each other. By 2014, XC2 and XC3 had moderate to high dispersion differences with every other stations. S36 also had differences compared to S42 and XC4, and XC4 was also different from both non-canyon stations.

## IV. Discussion

### 4.1 Temporal variability in DeSoto Canyon macrofauna and the adjacent slope

In this examination of temporal variability of Desoto Canyon macrofauna from 2012 – 2014, most community parameters showed no significant difference through time at a given station including total macrofaunal abundance, species richness, evenness, and abundance of most feeding guilds. Only a few parameters showed statistically significant changes at one or a few specific sites these include a decrease in abundance of total macrofauna at PCB06 and S36, and an increase in the abundance of deposit feeders decreased at PCB06, and in the abundance of chemosymbiotic species at XC3 between 2012 and 2014. Community structure, though not statistically significant, did shift over time within each station though, indicating some temporal variability in community structure, with a consistently larger change within each site from 2012 to 2013 than there was from 2013 to 2014. These somewhat low levels of change suggest that DeSoto Canyon communities were largely stable over the 3-year time period.

The lack of pronounced interannual variability in abundance at most stations is consistent with studies of non-canyon central NGOM polychaete assemblages spanning the continental shelf to the slope, from 1983 – 1984, in which Qu et al. (2017) and Reuscher and Shirley (2017) found relatively minor interannual variability in abundance, present in only a subset of the total sites investigated and often with the peak attributed to seasonal changes.

These results are inconsistent however, with other studies across the broader NGOM, that indicated temporal changes in abundance. For example, Pequegnat et al (1990) were the first to publish data to suggest interannual patterns in abundance among western, central, and eastern regions of the NGOM. From a later broad survey of the NGOM, Montagna et al. (2020) showed a significant 36% decrease in macrofaunal abundance from 2000 to 2001. Additionally, northwestern GOM macrofauna in a timespan close to the present study, 2010 – 2012, increased in abundance (Salcedo et al., 2017).

Inter-year investigations of diversity were not part of the focus of previous studies examining NGOM benthic temporal variability but community structure has been. Central NGOM continental slope polychaete assemblages showed large trajectory change in NMDS plots in 1983-1984, but when statistically tested, differences were minor, especially for sites deeper than 845 – 2540 m (Reuscher and Shirley, 2017). This is in keeping with the present study as the sites sampled in the DeSoto Canyon exhibited trajectories indicative of change and fall into the same depth range.

The short time period of the current study may have missed more gradual longer-term changes that can occur in benthic communities. Studies across decadal time scales may be needed to capture these patterns, but may also reveal long term stability in diversity in benthic macrofauna. For example Qu et al. (2017) and Reuscher and Shirley (2017) found polychaete assemblages variability between 1983 and 1984 though high, was not outside the range of variation from 2000 – 2001 at the same sites. Similarly Qu et al. (2017) observed diversity was similar for polychaetes between 1983 – 1985 and 2000 – 2002 on the western and central NGOM slope. Wei and Rowe (2019) observed the same diversity pattern for NGOM macrofauna as a whole across the same time period. Variability in community structure has less straightforward patterns in the long term. Distinct decadal separation in community structure was observed for upper continental slope polychaete assemblages (Qu et al., 2017) but not assemblages on the lower slope (Reuscher and Shirley, 2017).

Another possibility for long-term patterns is that there may be minimal long-term trajectories in communities that are punctuated by abrupt change brought on by stochastic events. In longer term time-series, polychaete assemblages in the northeast Atlantic, for example, displayed little change in density in the 5-6 years before the *‘Amperima’* event, wherein a suspected food pulse caused a rise in opportunistic cirratulids, spionids, and opheliids and also generated augmented numbers of deposit feeders and predators during and in the 2-3 years afterwards (Soto et al. 2010). In two long-term time-series of northern Pacific (1991 – 2005) and Atlantic (1991 – 1999) polychaetes, the greatest change in rank abundance of family and functional group did not occur until 1998, seven years after monitoring had started (Laguionie-Marchais et al. 2013). Temporal changes in community structure were also observed in that study as the 1998 time point tended to form a singular cluster, indicating some perturbation that year but no explanatory environmental factor could be identified (Laguionie-Marchais et al. 2013). Thus, greater temporal variability within the DeSoto Canyon cannot be ruled out as further research on DeSoto Canyon macrobenthic temporal dynamics over longer time scales may reveal patterns that are not apparent in a three-year temporal study.

### 4.2 Is there any evidence of the influence of the DwH oil spill on DeSoto Canyon macrofauna?

Patterns of macrobenthic oil spill response and recovery have primarily been investigated in shallow, coastal environments where recovery is a product of time and distance from the contamination source. After inundation by hydrocarbons, direct mortality can be observed, then rapid recolonization (months to a year) of opportunistic taxa occurs followed by a stabilization in abundance and richness (Berge 1990; Lu and Wu 2006). For macrofauna, this means initial increases in abundance driven by a dominance of deposit feeders, like polychaetes, and a drop in sensitive taxa such as crustaceans (especially amphipods) and molluscs (for an review of effects, see Suchanek (1993) and Peterson et al. (1996)). Large shifts in community composition would then result in a detectable alteration in community structure. Offshore GOM oil production impact assessments and reviews have generally corroborated the same effect, concluding continued industrial processes foster higher community dispersion, augmented abundances of deposit feeding polychaetes (driven by increased organic loading or resource availability), and are detrimental to sensitive crustaceans such as ostracods and amphipods (Montagna and Harper 1996; Peterson et al. 1996; Hernández-Arana et al. 2005). In terms of specific metrics of oil impact, more impacted sites and time points would be expected to have higher BPA ratios and higher dispersion (Andrade and Renaud, 2011). Among the DwH macrofaunal indicators, a greater proportion of tolerant and possibly tolerant groups along with lower relative abundances of sensitive and possibly sensitive taxa would be expected at impacted sites.

Schwing and Castillo (2020) posited oil spill recovery takes 3 – 5 years for GOM infauna based on evidence from foraminiferan abundance and recovery rates from the Ixtoc I oil spill in 1979 and the DwH. Sampling for the current study began two years after the initial spill, thus based on the patterns outlined above, if oil reached the canyon, sampling likely missed the initial phases of disturbance and the very start of the recovery process. It might be predicted that any impacted sites would then be in the phase of decline and stabilization in abundance, increasing diversity, and altered community structure through time. Year-to-year, only PCB06 and S36 exhibited any evidence of the expected impact patterns, showing peaks in abundance in 2012 decreasing into 2014. This lack of evidence in community patterns at most sites matches the lack of evidence for spill influence based on the oil spill metrics. Functional group abundance was only elevated for PCB06 in 2012 for deposit feeders and for chemosymbiotic taxa at XC3 in 2014. The increase in chemosymbiotic taxa was driven by an increase in thyasirid bivalves which are tolerant to DwH contamination (Washburn et al., 2016), and may suggest an increased influence at this site in 2014 compared to 2012. Deposit feeders increased at XC3 in 2014 also, which could support this inference, however the increase was not significant. There was a significant decrease in potentially tolerant species at XC3 in the same time period though, which contradicts the potential for increased oil influence at this site through time. Also, the increase in thyasirids at XC3 did not correspond with an increase in tolerant taxa at the same site over the study period, but XC3 had a higher proportion of tolerant taxa than any other site. For other stations, DwH indicator taxa proportions largely did not change through time except for a significant increase in potentially tolerant groups at XC2. Community dispersion was consistently high for XC4, but most other sites had dispersion comparable to or lower than non-canyon control stations.

Site-to-site, the number of differences found between macrofaunal communities among the oil-influence metrics were not high. Often, when differences were found, they were not detected by the final year of the sampling program. Sites with confirmed hydrocarbon deposition, PCB06, XC2, XC3, and XC4, most frequently exhibited greater potential indicators of impact compared to other canyon sites or adjacent slope sites. XC3 contained the most evidence for impact, but the evidence was stronger in 2014 rather than 2012. Chemosymbiotic feeders there in 2014 had elevated numbers compared to all other canyon stations and time points and, among the DwH macrofaunal indicators, contained greater proportions of tolerant and possibly tolerant taxa and low amounts of sensitive taxa compared to other canyon sites, especially in 2012. XC2 is the only other site to show low levels of sensitive taxa in both sampling years. XC4 showed the highest levels of dispersion (i.e., community stress) in all years. Finally, disparities among community dispersion were most frequent for XC2 and XC3 and persisted until the final year of sampling.

Taken together these metrics of oil influence indicate that the evidence for impacts to any of the stations was neither consistent among indicators nor overwhelming on a year-to-year or site-to-site basis. If any stations were still being affected by 2014, stations XC2, XC3, and XC4 had the greatest potential effect followed by PCB06. The depths of PCB06, XC2, and XC3, are consistent with the hydrocarbon plume depths (~1000 and 1400 m) reported by Hollander et al. (2012). The increased effects at XC3 in 2014 (plus high dispersion in 2012) plus the high dispersion at XC4 could potentially be explained by the turbidity-based downslope transport of hydrocarbons that Romero et al. (2020) proposed.

Severe infaunal impact first documented in close proximity and through time for the DwH (Qu et al. 2015; Montagna et al. 2016; Reuscher et al. 2017; Washburn et al. 2017) indicates the distance, low level of deposition, and time of sampling likely mitigated the overall effect observed for the DeSoto Canyon. Indeed, DeSoto Canyon stations are in the areas of the least petrocarbon deposition (Fig 1). Canyon sites are also located in regions of low impact according to predicted infaunal impact maps devised by Montagna et al (2013) and Reuscher et al. (2020) (Fig A1A and B). Chemical evidence would also suggest that impact to DeSoto Canyon macrofauna would not have been severe. For at least three of the sites in our target study area, XC2, PCB06, and XC3; total PAHs were elevated in 2010 (up to 524 ng g^−1^, 329 ng g^−1^, and 190 ng g^−1^ respectively) and 2011 (up to 99 ng g^−1^, 171 ng g^−1^, 373 ng g^−1^, respectively) following the DwH (Romero et al. 2015). Deposition levels were not measured or reported for XC4, but relatively heavy deposition can be inferred from the radiocarbon data presented in Figure 1B. These levels are generally greater than previous surveys of the canyon where background PAHs were observed to be 114 ± 47 ng g^−1^ (Rowe and Kennicutt 2009) but lower than the typical range of urbanized and industrialized coastal regions worldwide which span ~1200 – 11000 ng g^−1^ (Qiao et al. 2006; El Nemr et al. 2013). Further, concentrations reported by Romero et al. (2015) are also considered low to moderate pollution levels based on the proposed range of potential impact for aliphatics (10 – 100 μg g^−1^) and PAHs (100 – 1100 ng g^−1^) (Baumard et al. 1998; Commendatore et al. 2012). Observed PAH levels at these sites also fall below the “low limit” of PAH concentrations thought to be biologically toxic in shallow sediments (4402 ng g^−1^) (Long et al. 1995). Further, in an effort to establish sediment quality benchmarks for evaluating risks of oil-related impact to the deep-sea benthos from the DwH, Balthis et al. (2017) derived a low probability of impact to macrofauna (< 20%) when total PAH concentrations were lower than 4000 ng g^−1^ (ppm) and high (> 80%) when greater than 24000 ng g^−1^. Thus, it can be concluded the level of hydrocarbon influx into the DeSoto Canyon would not have severely impacted resident biota and this conclusion is largely corroborated by the scant effects observed in the macrofauna communities of the present study.

## IV. Conclusions

DeSoto Canyon macrofauna exhibited limited amounts of temporal change in abundance, diversity, and community structure through the study period. Similarly, attempts to relate observed trends to the DwH using established faunal oil-impact and disturbance metrics indicated little to no evidence of impact. Sites with confirmed hydrocarbon deposition, XC2, XC3, and XC4, had minor evidence of potential impacts, but results were not consistent across metrics or years. The lack of temporal change and response to the DwH is likely a product of distance from the wellhead (~40 – 185 km east/southeast), time of sampling, and the general low deposition of DWH hydrocarbons within the canyon.

