## Supplemental Tables and Figures for "Interannual temporal patterns of DeSoto Canyon macrofauna and evaluation of influence from the Deepwater Horizon"

Table A1. Summary of the PERMANOVA results for the feeding guild abundances. Sample size n = 3 for all groups in pairwise comparisons except for NT800 in 2012 when n = 2. Bolded values indicate significant differences for 2012 and 2013 when α is below 0.005, 2014 when α is below 0.0024.

| | | Carnivore | | | Deposit | | | Omnivore | | | Suspension feeder | | | Chemosymbiotic | | |
|---|---|---|---|---|---|---|---|---|---|---|---|---|---|---|---|---|
| | df | SS | Pseudo-F | *P* | SS | Pseudo-F | *P* | SS | Pseudo-F | *P* | SS | Pseudo-F | *P* | SS | Pseudo-F | *P* |
| Time Point | 2 | 87.085 | 0.89508 | 0.415 | 71.892 | 1.454 | 0.244 | 129.92 | 0.6445 | 0.576 | 860.1 | 1.1643 | 0.347 | 214.8 | 0.7025 | 0.537 |
| Station | 6 | 7391.1 | 25.322 | **0.001** | 2861 | 19.29 | **0.001** | 7445.5 | 12.312 | **0.001** | 3217 | 1.4517 | 0.15 | 7228.8 | 8.4258 | **0.001** |
| Time Point x Station | 8 | 250.64 | 0.64402 | 0.739 | 322.3 | 1.6294 | 0.149 | 421.65 | 0.5229 | 0.887 | 2117.1 | 0.7165 | 0.873 | 2863.6 | 2.3414 | **0.03** |
| Residual | 33 | 1605.3 | | 815.91 | | | 3326.1 | | | 12189 | | | 5045 | | | |
| Total | 49 | 9491 | | 4390.7 | | | 11410 | | | 19031 | | | 16131 | | | |

| | Carnivore | | | Deposit | | | Omnivore | | | Suspension | | | Chemosymbiotic | | |
|---|---|---|---|---|---|---|---|---|---|---|---|---|---|---|---|
| | 2012 | 2013 | 2014 | 2012 | 2013 | 2014 | 2012 | 2013 | 2014 | 2012 | 2013 | 2014 | 2012 | 2013 | 2014 |
| Groups | *P* | *P* | *P* | *P* | *P* | *P* | *P* | *P* | *P* | *P* | *P* | *P* | *P* | *P* | *P* |
| S42, NT800 | - | 0.798 | 0.745 | - | 0.546 | 0.28 | - | 0.499 | 0.119 | - | 0.083 | 0.784 | - | 0.528 | 0.354 |
| S42, XC2 | - | - | **0.001** | - | - | **0.002** | - | - | 0.022 | - | - | 0.108 | - | - | 0.449 |
| S42, XC3 | - | - | 0.047 | - | - | 0.011 | - | - | 0.063 | - | - | 0.55 | - | - | 0.035 |
| NT800, XC2 | - | - | 0.008 | - | - | 0.01 | - | - | 0.162 | - | - | 0.355 | - | - | 0.888 |
| NT800, XC3 | - | - | 0.317 | - | - | 0.033 | - | - | 0.265 | - | - | 0.83 | - | - | 0.008 |
| XC2, XC3 | 0.028 | - | **0.002** | 0.373 | - | 0.276 | 0.03 | - | 0.595 | 0.686 | - | 0.17 | 0.829 | - | 0.011 |
| PCB06, S36 | 0.086 | 0.041 | 0.01 | **0.003** | 0.324 | 0.237 | 0.016 | 0.739 | 0.412 | 0.482 | 0.75 | 0.763 | 0.051 | 0.205 | 0.036 |
| PCB06, XC4 | **0.001** | 0.005 | 0.015 | 0.014 | 0.005 | 0.003 | 0.093 | 0.046 | 0.074 | 0.057 | 0.063 | 0.365 | 0.038 | 0.043 | 0.045 |
| PCB06, XC2 | 0.928 | - | 0.01 | 0.24 | - | 0.034 | 0.492 | - | 0.43 | 0.315 | - | 0.517 | 0.553 | - | 0.205 |
| PCB06, XC3 | 0.035 | - | 0.004 | 0.2 | - | 0.183 | 0.024 | - | 0.349 | 0.365 | - | 0.553 | 0.711 | - | 0.029 |
| PCB06, S42 | - | 0.097 | **0.001** | - | 0.113 | 0.066 | - | 0.406 | 0.642 | - | 0.013 | 0.332 | - | 0.496 | 0.659 |
| PCB06, NT800 | - | 0.132 | 0.024 | - | 0.278 | 0.284 | - | 0.065 | 0.735 | - | 0.24 | 0.611 | - | 0.296 | 0.133 |
| S36, XC4 | **0.004** | 0.085 | 0.126 | 0.09 | **0.003** | 0.061 | 0.033 | 0.062 | 0.036 | 0.026 | 0.433 | 0.351 | Neg | 0.41 | Neg |
| S36, XC2 | 0.061 | - | 0.003 | 0.065 | - | 0.022 | 0.012 | - | 0.611 | 0.499 | - | 0.208 | 0.043 | - | 0.776 |
| S36, XC3 | 0.349 | - | 0.18 | 0.89 | - | 0.055 | 0.768 | - | 0.835 | 0.524 | - | 0.724 | 0.048 | - | **0.002** |
| S36, S42 | - | 0.559 | 0.639 | - | 0.352 | 0.865 | - | 0.689 | 0.03 | - | 0.274 | 0.359 | - | 0.08 | 0.229 |
| S36, NT800 | - | 0.847 | 0.932 | - | 0.683 | 0.585 | - | 0.329 | 0.163 | - | 0.802 | 0.719 | - | 0.085 | 0.867 |
| XC4, XC2 | **0.001** | - | 0.005 | 0.027 | - | **0.001** | 0.076 | - | 0.033 | 0.071 | - | 0.338 | 0.034 | - | 0.766 |
| XC4, XC3 | 0.012 | - | 0.042 | 0.158 | - | **0.001** | 0.027 | - | 0.034 | 0.518 | - | 0.4 | 0.041 | - | **0.002** |
| XC4, S42 | - | 0.034 | 0.083 | - | **0.001** | 0.003 | - | 0.096 | 0.078 | - | 0.434 | 0.433 | - | 0.011 | 0.211 |
| XC4, NT800 | - | 0.131 | 0.131 | - | **0.002** | 0.006 | - | 0.224 | 0.047 | - | 0.315 | 0.489 | - | 0.015 | 0.843 |

Table A1 continued

| | Carnivore | | | Deposit | | | Omnivore | | | Suspension | | | Chemosymbiotic | | |
|---|---|---|---|---|---|---|---|---|---|---|---|---|---|---|---|
| | 2012 | 2013 | 2014 | 2012 | 2013 | 2014 | 2012 | 2013 | 2014 | 2012 | 2013 | 2014 | 2012 | 2013 | 2014 |
| Groups | t | t | t | t | t | t | t | t | t | t | t | t | t | t | t |
| S42, NT800 | - | 0.2969 | 0.32351 | - | 0.68605 | 1.288 | - | 0.71462 | 1.9468 | - | 2.5098 | 0.37622 | - | 0.72555 | 1.0313 |
| S42, XC2 | - | - | 13.839 | - | - | 6.7212 | - | - | 3.6863 | - | - | 2.0119 | - | - | 0.8339 |
| S42, XC3 | - | - | 2.9198 | - | - | 4.8479 | - | - | 2.3004 | - | - | 0.6993 | - | - | 2.8024 |
| NT800, XC2 | - | - | 4.9191 | - | - | 4.847 | - | - | 1.7558 | - | - | 1.0565 | - | - | 0.15852 |
| NT800, XC3 | - | - | 1.144 | - | - | 3.3085 | - | - | 1.2665 | - | - | 0.35358 | - | - | 4.0578 |
| XC2, XC3 | 3.4554 | - | 9.1197 | 0.99908 | - | 1.2612 | 3.7184 | - | 0.59672 | 0.44587 | - | 1.6788 | 0.27414 | - | 3.6112 |
| PCB06, S36 | 2.2546 | 3.1214 | 4.8989 | 9.6524 | 1.0932 | 1.3995 | 3.9359 | 0.38756 | 0.97193 | 0.79117 | 0.38679 | 0.3463 | 2.5647 | 1.4455 | 2.9634 |
| PCB06, XC4 | 6.5664 | 5.8176 | 4.1935 | 3.9794 | 6.0605 | 6.3958 | 2.1452 | 2.7896 | 2.1805 | 2.815 | 2.4428 | 1.0176 | 2.5647 | 2.6815 | 2.9634 |
| PCB06, XC2 | 0.10812 | - | 4.7136 | 1.3398 | - | 2.9477 | 0.72974 | - | 0.82967 | 1.1763 | - | 0.72895 | 0.6752 | - | 1.4677 |
| PCB06, XC3 | 3.1995 | - | 7.5095 | 1.6135 | - | 1.717 | 3.3516 | - | 0.98609 | 1.0386 | - | 0.62025 | 0.41525 | - | 3.2944 |
| PCB06, S42 | - | 2.2859 | 19.415 | - | 2.0281 | 2.4399 | - | 0.98712 | 0.4938 | - | 3.8173 | 1.151 | - | 0.7669 | 0.50522 |
| PCB06, NT800 | - | 2.0861 | 3.5237 | - | 1.3065 | 1.2468 | - | 2.7753 | 0.37287 | - | 1.4871 | 0.58434 | - | 1.2855 | 1.7513 |
| S36, XC4 | 5.6219 | 2.331 | 1.9084 | 2.2892 | 7.152 | 2.6086 | 3.0734 | 2.3913 | 3.2512 | 3.4654 | 0.87454 | 1.0192 | Neg | 0.91821 | Neg |
| S36, XC2 | 2.4648 | - | 6.5354 | 2.5894 | - | 3.6156 | 5.0252 | - | 0.58061 | 0.73942 | - | 1.4881 | 3.0389 | - | 0.36426 |
| S36, XC3 | 1.1363 | - | 1.651 | 0.1666 | - | 2.7245 | 0.32586 | - | 0.26288 | 0.74513 | - | 0.39355 | 2.7241 | - | 7.9155 |
| S36, S42 | - | 0.66184 | 0.56813 | - | 1.1277 | 0.19713 | - | 0.44394 | 3.2158 | - | 1.2921 | 1.0524 | - | 2.2799 | 1.4605 |
| S36, NT800 | - | 0.21323 | 0.1105 | - | 0.46946 | 0.58079 | - | 1.1287 | 1.6851 | - | 0.34004 | 0.52051 | - | 2.5479 | 0.25235 |
| XC4, XC2 | 6.8769 | - | 4.9962 | 3.1617 | - | 12.827 | 2.37 | - | 3.217 | 2.3256 | - | 1.1411 | 3.0389 | - | 0.36426 |
| XC4, XC3 | 4.8888 | - | 2.8242 | 1.5798 | - | 9.841 | 3.0018 | - | 3.1031 | 0.73793 | - | 0.97821 | 2.7241 | - | 7.9155 |
| XC4, S42 | - | 2.9585 | 2.3512 | - | 9.066 | 5.6238 | - | 2.1756 | 2.3821 | - | 0.86184 | 0.91543 | - | 3.9111 | 1.4605 |
| XC4, NT800 | - | 1.9683 | 1.8364 | - | 11.322 | 6.1099 | - | 1.5352 | 2.8899 | - | 1.1973 | 0.85393 | - | 4.5886 | 0.25235 |

Table A2. Summary of PERMANOVA test of Deepwater Horizon macrofaunal indicator proportions. Sample size n = 3 for all groups in pairwise comparisons except for NT800 in 2012 where n =2. Bolded values indicate significant differences for 2012 and 2013 when α is below 0.005, 2014 when α is below 0.0024.

| | | Cosmopolitan | | | Tolerant | | | Possibly tolerant | | | Possibly sensitive | | | Sensitive | | |
|---|---|---|---|---|---|---|---|---|---|---|---|---|---|---|---|---|
| | df | SS | Pseudo-F | *P* | SS | Pseudo-F | *P* | SS | Pseudo-F | *P* | SS | Pseudo-F | *P* | SS | Pseudo-F | *P* |
| Time Point | 2 | 1.0225 | 0.91342 | 0.424 | 105.03 | 2.1075 | 0.135 | 4.69E-02 | 0.21704 | 0.786 | 24.899 | 0.34496 | 0.72 | 39.798 | 1.5825 | 0.217 |
| Site | 6 | 53.009 | 15.785 | **0.001** | 3509.3 | 23.471 | **0.001** | 3.3309 | 5.1431 | **0.002** | 408.58 | 5.6608 | **0.001** | 1699.4 | 22.524 | **0.001** |
| Time Point x Site | 8 | 4.8096 | 1.0741 | 0.432 | 198.77 | 0.99708 | 0.461 | 1.5565 | 1.8025 | 0.115 | 142.96 | 1.9806 | 0.08 | 26.49 | 2.2515 | **0.027** |
| Residual | 33 | 18.47 | | | 822.33 | | | 3.562 | | | 72.178 | | | 414.95 | | |
| Total | 49 | 77.268 | | | 4926.4 | | | 8.7291 | | | | | | 2702.8 | | |

| | Cosmopolitan | | | Tolerant | | | Possibly tolerant | | | Possibly sensitive | | | Sensitive | | |
|---|---|---|---|---|---|---|---|---|---|---|---|---|---|---|---|
| | 2012 | 2013 | 2014 | 2012 | 2013 | 2014 | 2012 | 2013 | 2014 | 2012 | 2013 | 2014 | 2012 | 2013 | 2014 |
| Groups | *P* | *P* | *P* | *P* | *P* | *P* | *P* | *P* | *P* | *P* | *P* | *P* | *P* | *P* | *P* |
| S42, NT800 | - | **0.004** | 0.528 | - | 0.792 | 0.311 | - | 0.031 | 0.592 | - | 0.828 | 0.449 | - | 0.233 | 0.182 |
| S42, XC2 | - | - | 0.21 | - | - | 0.908 | - | - | 0.088 | - | - | 0.206 | - | - | 0.01 |
| S42, XC3 | - | - | 0.185 | - | - | 0.004 | - | - | 0.699 | - | - | 0.106 | - | - | 0.117 |
| NT800, XC2 | - | - | 0.183 | - | - | 0.181 | - | - | 0.224 | - | - | 0.097 | - | - | **0.001** |
| NT800, XC3 | - | - | 0.868 | - | - | 0.004 | - | - | 0.35 | - | - | 0.625 | - | - | 0.052 |
| XC2, XC3 | **0.002** | - | 0.046 | 0.095 | - | 0.003 | 0.231 | - | 0.01 | 0.925 | - | 0.003 | 0.025 | - | 0.44 |
| PCB06, S36 | 0.498 | 0.866 | 0.022 | 0.319 | 0.348 | 0.916 | 0.31 | 0.993 | 0.043 | 0.532 | 0.006 | 0.098 | 0.266 | 0.115 | 0.032 |
| PCB06, XC4 | 0.038 | 0.015 | 0.134 | 0.172 | 0.016 | 0.38 | 0.032 | 0.118 | 0.006 | 0.074 | 0.298 | 0.302 | 0.045 | 0.101 | 0.08 |
| PCB06, XC2 | 0.015 | - | 0.102 | 0.185 | - | 0.405 | 0.248 | - | 0.13 | 0.291 | - | 0.205 | 0.2 | - | 0.016 |
| PCB06, XC3 | 0.514 | - | 0.163 | 0.009 | - | 0.03 | 0.78 | - | 0.062 | 0.669 | - | 0.092 | 0.011 | - | 0.364 |
| PCB06, S42 | - | 0.009 | 0.697 | - | 0.071 | 0.437 | - | 0.01 | 0.336 | - | 0.654 | 0.972 | - | 0.334 | 0.212 |
| PCB06, NT800 | - | 0.769 | 0.596 | - | 0.07 | 0.161 | - | 0.674 | 0.766 | - | 0.568 | 0.451 | - | 0.187 | 0.028 |
| S36, XC4 | 0.014 | 0.02 | 0.481 | 0.024 | 0.035 | 0.256 | 0.408 | 0.335 | 0.003 | 0.198 | 0.047 | 0.813 | 0.071 | 0.806 | 0.743 |
| S36, XC2 | 0.008 | - | 0.025 | 0.474 | - | 0.087 | 0.827 | - | 0.005 | 0.334 | - | 0.019 | 0.026 | - | **0.002** |
| S36, XC3 | 0.04 | - | 0.585 | **0.001** | - | 0.007 | 0.311 | - | 0.873 | 0.454 | - | 0.548 | **0.001** | - | 0.051 |
| S36, S42 | - | 0.019 | 0.073 | - | 0.013 | 0.18 | - | 0.219 | 0.738 | - | 0.007 | 0.112 | - | 0.191 | 0.2 |
| S36, NT800 | - | 0.756 | 0.678 | - | 0.027 | 0.051 | - | 0.879 | 0.353 | - | 0.019 | 0.489 | - | 0.697 | 0.671 |
| XC4, XC2 | 0.007 | - | 0.051 | 0.023 | - | 0.062 | 0.096 | - | **0.001** | 0.045 | - | 0.071 | 0.008 | - | 0.007 |
| XC4, XC3 | 0.042 | - | 0.322 | 0.049 | - | 0.083 | **0.003** | - | 0.024 | 0.078 | - | 0.869 | **0.001** | - | 0.067 |
| XC4, S42 | - | **0.001** | 0.113 | - | **0.004** | 0.094 | - | 0.709 | 0.131 | - | 0.214 | 0.256 | - | 0.213 | 0.286 |
| XC4, NT800 | - | 0.006 | 0.406 | - | **0.003** | 0.034 | - | 0.164 | 0.068 | - | 0.308 | 0.696 | - | 0.585 | 0.643 |

Table A2 continued.

| | Cosmopolitan | | | Tolerant | | | Possibly tolerant | | | Possibly sensitive | | | Sensitive | | |
|---|---|---|---|---|---|---|---|---|---|---|---|---|---|---|---|
| | 2012 | 2013 | 2014 | 2012 | 2013 | 2014 | 2012 | 2013 | 2014 | 2012 | 2013 | 2014 | 2012 | 2013 | 2014 |
| Groups | t | t | t | t | t | t | t | t | t | t | t | t | t | t | t |
| S42, NT800 | - | 7.1211 | 0.73724 | - | 0.27449 | 1.2339 | - | 3.8353 | 0.56391 | - | 0.26499 | 0.83871 | - | 1.4462 | 1.6147 |
| S42, XC2 | - | - | 1.5164 | - | - | 0.11015 | - | - | 2.4167 | - | - | 1.4851 | - | - | 5.832 |
| S42, XC3 | - | - | 1.6196 | - | - | 5.5833 | - | - | 0.42339 | - | - | 2.1372 | - | - | 1.919 |
| NT800, XC2 | - | - | 1.7034 | - | - | 1.5591 | - | - | 1.4305 | - | - | 2.0456 | - | - | 11.861 |
| NT800, XC3 | - | - | 0.16994 | - | - | 5.9755 | - | - | 1.0696 | - | - | 0.56234 | - | - | 2.7896 |
| XC2, XC3 | 7.3925 | - | 2.8934 | 6.1445 | - | 6.7572 | 1.435 | - | 5.0172 | 0.12574 | - | 5.8066 | 3.1882 | - | 0.89733 |
| PCB06, S36 | 0.69589 | 0.15737 | 3.5815 | 1.1518 | 1.1057 | 0.13206 | 1.1323 | 0.013835 | 2.7798 | 0.68601 | 7.033 | 1.9901 | 1.3335 | 2.0486 | 3.3243 |
| PCB06, XC4 | 2.9815 | 4.2377 | 1.9345 | 1.719 | 3.925 | 0.98993 | 3.1566 | 2.0557 | 6.1909 | 2.4549 | 1.2607 | 1.2463 | 2.7378 | 2.0238 | 2.2683 |
| PCB06, XC2 | 3.8542 | - | 2.1241 | 1.5755 | - | 0.94255 | 1.2578 | - | 1.9503 | 1.1698 | - | 1.5565 | 1.4676 | - | 3.8696 |
| PCB06, XC3 | 0.75542 | - | 1.768 | 4.4572 | - | 3.2071 | 0.26815 | - | 2.6225 | 0.47767 | - | 2.0782 | 4.704 | - | 1.066 |
| PCB06, S42 | - | 4.935 | 0.44319 | - | 2.496 | 0.89108 | - | 5.1514 | 1.1463 | - | 0.47058 | 0.053222 | - | 1.1963 | 1.4594 |
| PCB06, NT800 | - | 0.31054 | 0.58196 | - | 2.8155 | 1.719 | - | 0.4435 | 0.33419 | - | 0.62711 | 0.79856 | - | 1.6901 | 3.4302 |
| S36, XC4 | 4.1591 | 3.5156 | 0.83798 | 3.5228 | 2.8547 | 1.2855 | 0.88549 | 1.0869 | 5.7123 | 1.5145 | 2.8576 | 0.26979 | 2.4767 | 0.2523 | 0.3408 |
| S36, XC2 | 4.5118 | - | 3.6089 | 0.82716 | - | 2.3443 | 0.24494 | - | 5.5642 | 1.1288 | - | 3.9707 | 3.2714 | - | 9.8345 |
| S36, XC3 | 2.84 | - | 0.64547 | 8.3732 | - | 4.893 | 1.1172 | - | 0.19316 | 0.82566 | - | 0.62912 | 8.0896 | - | 2.8386 |
| S36, S42 | - | 3.8185 | 2.4615 | - | 3.605 | 1.6466 | - | 1.4616 | 0.3483 | - | 6.0143 | 2.0361 | - | 1.6463 | 1.6872 |
| S36, NT800 | - | 0.39055 | 0.46781 | - | 4.0919 | 2.713 | - | 0.171 | 1.0229 | - | 4.4391 | 0.80142 | - | 0.47065 | 0.4291 |
| XC4, XC2 | 6.6893 | - | 2.8653 | 3.339 | - | 2.4828 | 2.01 | - | 9.9192 | 2.7677 | - | 2.4264 | 4.7085 | - | 4.9844 |
| XC4, XC3 | 3.1852 | - | 1.0832 | 2.8204 | - | 2.2905 | 5.2825 | - | 3.6463 | 2.2706 | - | 0.25748 | 9.7557 | - | 2.5242 |
| XC4, S42 | - | 14.275 | 1.9785 | - | 6.4175 | 2.2305 | - | 0.39734 | 1.962 | - | 1.4457 | 1.2827 | - | 1.5851 | 1.2741 |
| XC4, NT800 | - | 6.6029 | 0.94467 | - | 8.7495 | 2.9867 | - | 1.7881 | 2.4216 | - | 1.2607 | 0.43632 | - | 0.64718 | 0.51872 |

Figure A1) Time-series sites in relation to benthic DwH faunal impact interpolation maps. A) Based on data from Montagna et al. (2013), the interpolated area of deep sea impact are based on PC1 station scores of infaunal (meiofauna and macrofauna) data covering 70,166 $km^2$. Orange (167 $km^2$) are moderately impacted and red (24 $km^2$) are severely impacted. High diversity and low chemical loads (yellow/green) are unimpacted. B) Based on data from Montagna et al. (2020) and contour colors follow Montagna et al. (2013).

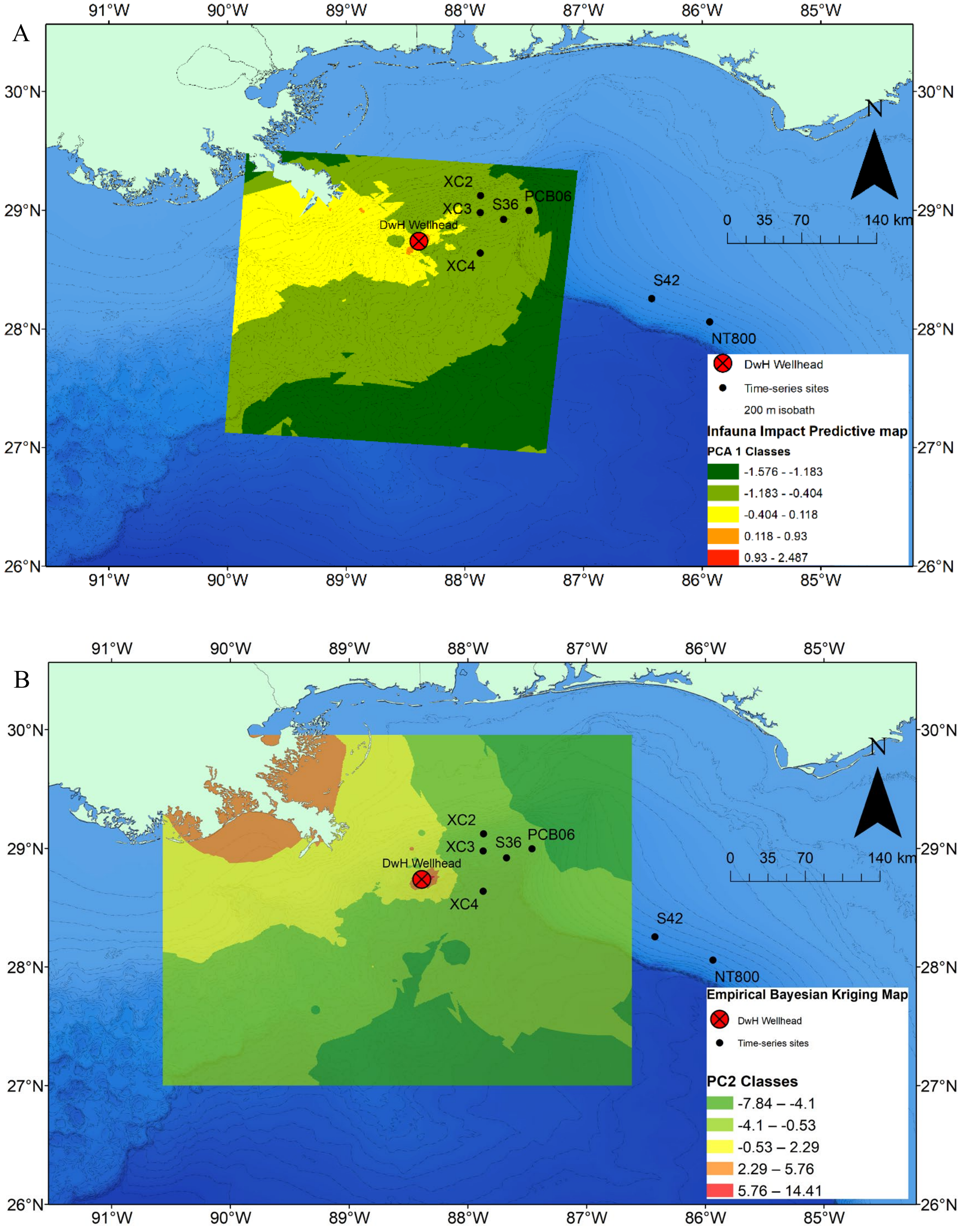
